# Mapping the Immune Landscape of Clear Cell Renal Cell Carcinoma by Single-Cell RNA-seq

**DOI:** 10.1101/824482

**Authors:** Ajaykumar Vishwakarma, Nicholas Bocherding, Michael S. Chimenti, Purshottam Vishwakarma, Kenneth Nepple, Aliasger Salem, Russell W. Jenkins, Weizhou Zhang, Yousef Zakharia

## Abstract

The immune cells within the tumor microenvironment are considered key determinants of response to cancer immunotherapy. Immune checkpoint blockade (ICB) has transformed the treatment of clear cell renal cell carcinoma (ccRCC), although the role of specific immune cell states remains unclear. To characterize the tumor microenvironment (TME) of ccRCC, we applied single-cell RNA sequencing (scRNA-seq) along with paired T cell receptor sequencing to map the transcriptomic heterogeneity of 24,904 individual CD45^+^ lymphoid and myeloid cells in matched tumor and blood from patients with ccRCC. We identified multiple distinct immune cell phenotypes for B and T (CD4 and CD8) lymphocytes, natural kill (NK) cells, and myeloid cells. Evaluation of T cell receptor (TCR) sequences revealed limited shared clonotypes between patients, whereas tumor-infiltrating T cell clonotypes were frequently found in peripheral blood, albeit in lower abundance. We further show that the circulating CD4^+^ T cell clonality is far less diverse than peripheral CD8^+^. Evaluation of myeloid subsets revealed unique gene programs defining monocytes, dendritic cells and tumor-associated macrophages. In summary, here we have leveraged scRNA-seq to refine our understanding of the relative abundance, diversity and complexity of the immune landscape of ccRCC. This report represents the first characterization of ccRCC immune landscape using scRNA-seq. With further characterization and functional validation, these findings may identify novel sub-populations of immune cells amenable to therapeutic intervention.

**One Sentence Summary:** Single-cell RNA-sequencing reveals unique lymphoid and myeloid gene programs with putative functions in clear cell renal cancer patients

## Introduction

ccRCC is the most common type of renal cell carcinoma arising from epithelial cells of the proximal tubule of the kidney, comprising more than 70% of all renal cancers (*1*). ccRCC represents an immune sensitive tumor type well known for early advances in systemic immunotherapy using T cell proliferation cytokine IL-2 and interferon-α2b therapy (*2*). Recent novel immunotherapies targeting T cell immune checkpoints as standard of care has transformed the treatment paradigm of ccRCC (*3, 4*). However, a substantial subset of renal cancer patients still do not respond to these therapies and many patients who initially do respond can eventually progress (*5, 6*). Cytotoxic CD8^+^ T cells are key effectors of the adaptive anti-tumor immune response (*7*) and the abundance of CD8^+^ T cells in solid cancers is generally associated with better survival in cancer patients (*8–10*). However, in RCC, tumor-infiltrating lymphocyte (TIL) abundance is inversely correlated with survival (*11–13*). Biomarker analysis from recent clinical trials comparing PD-1 blockade versus anti-angiogenic inhibitors and combination therapies in treatment-naïve ccRCC patients also demonstrated the inverse relationship between T cell infiltration and clinical outcomes (*14, 15*). Other abundant immune players in the ccRCC TME include monocytes, dendritic cells, and tumor-associated macrophages (*16*), which are now being harnessed for discovery of novel gene programs but remain far less studied than T cells.

Quantifying and inferring immune cell abundance from transcriptional analysis of primary or metastasized bulk tumor samples is inadequate to provide a clear picture of the immune cell types (*17, 18*). While these studies are suggestive, they lack single cell resolution for characterizing heterogeneous cell subpopulations that ultimately shape anti-tumor response, as has been demonstrated in breast cancer and melanoma (*19, 20*). Single cell methodologies including flow cytometry, immunohistochemistry and mass cytometry (*13, 16, 21*) have revealed immune cell states in ccRCC but only as discrete phenotypes when in vivo they typically display diverse spectrum of differentiation or activation states. Also, these methods require the use of antibody panels targeting known immune cell components, and by design are not capable of identifying novel cell states. scRNA-seq has enabled comprehensive characterization of heterogeneous lymphoid and myeloid immune cells in several cancers (*22–25*), providing an unbiased approach to profile cells and enable molecular classification of different subpopulations and identification of novel gene programs. Transcriptome mapping of T lymphocytes coupled with TCR sequencing allows additional measurement of clonal T cell response to cancer at an unprecedented depth (*26, 27*).

Here, we report the single cell RNA profiling of the immune landscape in ccRCC mapping a total 24,904 of immune cells (5’ gene expression and recombined V(D)J region of the T cell receptor) in matched samples of tumor and peripheral blood isolated from three treatment-naïve ccRCC patients. Analysis of CD45^+^ TILs revealed remarkable heterogeneity of CD8^+^ and CD4^+^ T cell sub-populations compared to peripheral blood. Pseudotime ordering of these sub-populations further revealed the gene program adaptations during their transition from peripheral bloods to tumor tissues. We characterized the clonal structure of T cells from tumors and paired peripheral blood specific to each patient. Analysis of myeloid cells revealed a complex mixture of pro- and anti-inflammatory polarized phenotypes across patients. This represents the first such report of the immune landscape of ccRCC using scRNA-seq.

## Results

### Single Immune Cell Transcriptome Data Workflow, Analysis and Cellular Identity of Purified 24,904 Primary Immune Cells in ccRCC

To define lymphoid and myeloid sub-populations in human ccRCC, we first obtained tumor and peripheral blood specimens of three treatment-naïve ccRCC patients. To establish relative abundance of immune cell types particularly within tumor; and between tumor and peripheral blood, we used conventional lymphoid and myeloid gates during flow cytometric sorting of single cells to pool the two broad cell types in different proportions (lymphoid: myeloid, 7:3) across the three-patients. The flow sorted lymphoid and myeloid cells were processed via the 10X scRNA-seq platform (Fig. 1A). The merged tumor and blood data from each patient was visualized for exploration via Loupe Cell Browser (Fig. S1A-C), and after filtering scRNA-seq gene expression data, a total of 25,688 cell transcriptomes at an average sequencing depth of 50,000 reads per cell were retained for further analysis across all the ccRCC patients.

**Fig. 1.**
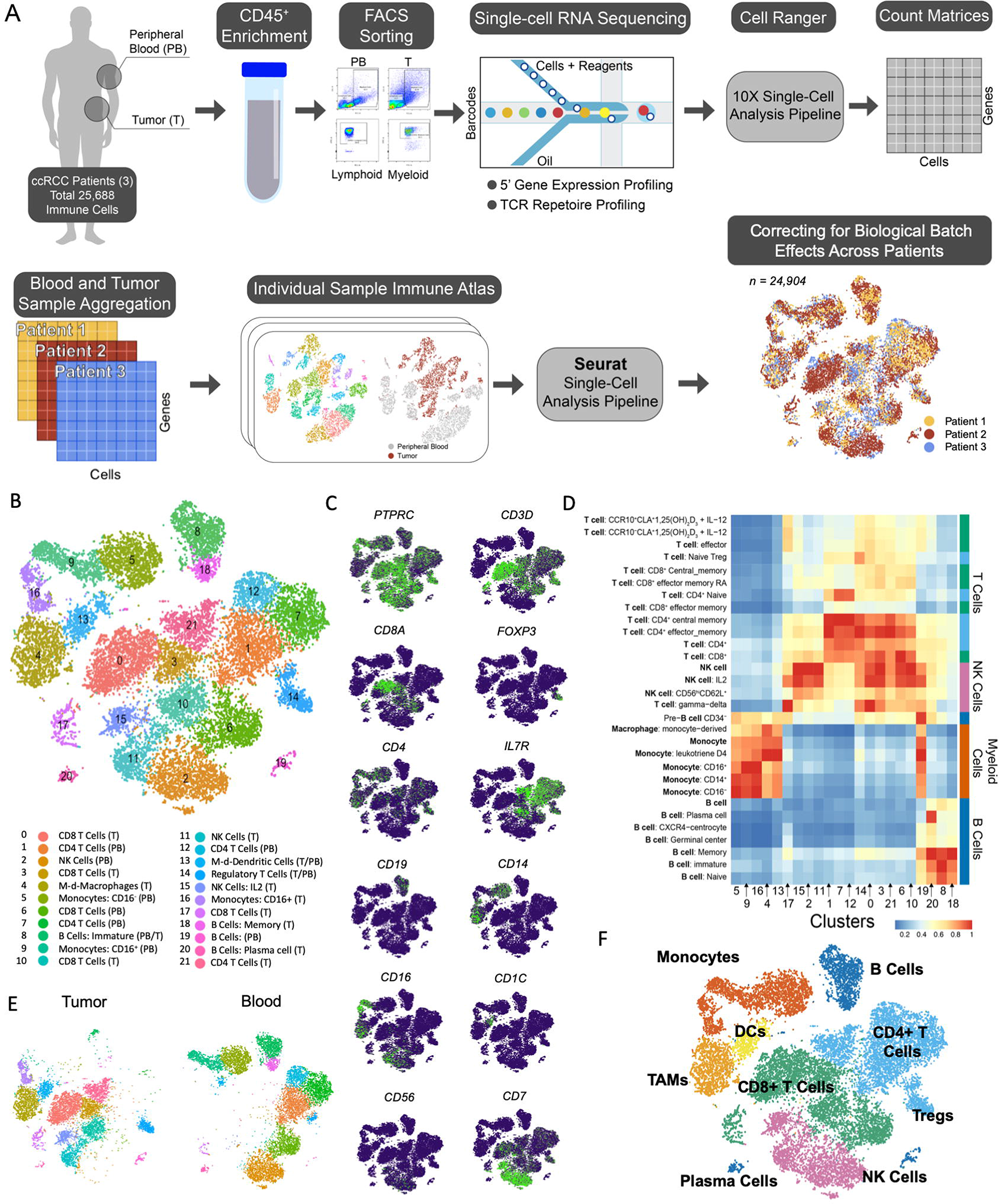
Single-Cell RNA sequencing Data Workflow, Analysis and Cellular Identity of Purified 25,688 Primary Immune Cells in ccRCC: **(A)** Workflow for single cell RNA-seq of isolated lymphoid and myeloid immune cell populations from ccRCC subjects and computational analysis. Single cell RNA-seq data across patients is normalized for library size and clustered using Seurat. After correcting for batch effects across individuals, cells are colored by patient **(B)** tSNE plot of complete immune cell atlas from 3 ccRCC tumors and paired peripheral blood. **(C)** Gene expression of canonical immune cell markers in each cluster **(D)** Spearman’s correlation of cluster expression co-related to Human Primary Cell Atlas (HPCA) dataset **(E)** tSNE plot of immune cell transcriptomes from patient tumor (n=3) and blood samples (n=3) (F) tSNE plot shows lymphoid and myeloid immune cell subsets in ccRCC. Cells colored by inferred immune cell type.

To obtain an unbiased systematic comparison across patients, we merged the tumor and blood data from all three-patients. Upon performing single integrated analysis on cells from all three patient, we observed optimized overlapping of clusters across the biological replicates, indicative of extensive similarities and therefore, we set out to define the immune sub-populations across the limited specimens. After merging Seurat normalized dataset, 24,904 cells remained across all three-patient’s paired tumor and blood. We began by clustering these cells using Seurat’s unsupervised clustering algorithm (*28*) and constructed 22 distinct clusters (0-21) represented by t-distributed stochastic neighbor embedding (tSNE) approach (Fig.1B). By inspection of cell type specific gene expression, we observed that each cluster defined a unique gene signature likely representative of distinct cell type. tSNE projections were able to capture majority of the anticipated lymphoid and myeloid cell-types expressing *PTPRC*, the gene encoding the antigen CD45 (Fig.1C). To confirm the identity of cells within each cluster, we examined the expression levels of canonical markers for the T cells (*CD3, CD8, CD4, FOXP3)*, B cells *(CD19)*, natural killer (NK) cells *(CD7*, *CD56)* and myeloid cell sub-types; monocytes *(CD14, FCGR3A)*, macrophages *(APOE*, *APOC1*, *MRC1)* and dendritic cells *(CD1c*, *FCERA)* (Fig.1C). The identity of single cells was further determined by correlating cluster mean expression of the immune cell signatures extracted from the human primary cell atlas (HPCA), a collection of Gene Expression Omnibus (GEO) datasets containing 713 microarray samples that were classified into 37 main cell types and further annotated to 157 sub-types (Fig.1D). Of note, there was a marked distinction in the expression profiles between tumor-infiltrating immune cells and peripheral blood mononuclear cells (PBMCs) with clear separation between tumor and blood (Fig.1E).

Using single-cell transcriptomes from paired blood and tumor specimens from three-patients with ccRCC, we identified major lymphoid and myeloid immune cell lineages (Fig. 1F). We annotated a total of 10 T cell clusters, 5 myeloid clusters, 3 NK cell and 4 B cell clusters. The complete differential gene expression results across all clusters are available (Table S1). Only subtle difference between cell type frequencies were observed amongst patients, with the exception of B cells (Table S1). Our strategy of single cell analysis performed on immune cells taking into consideration the frequencies of lymphoid and myeloid cells during flow sorting provided a powerful way to identify relative abundance between proportion of cell types and corresponding immune cell states. CD8^+^ T cells were the most abundant intra-tumoral CD3^+^ immune cell population (Fig. 1F), whereas blood-specific clusters demonstrated a higher fraction of CD4^+^ T cell infiltrates. Tumor-associated monocytes/macrophages (TAMs) constituted the largest intra-tumoral myeloid cell cluster in our data.

### CD8^+^ T Cells Exhibit Continuum of Distinct Transcriptional States within Tumor and Peripheral Blood

Cytotoxic CD8^+^ T lymphocytes are recognized as the key effectors of adaptive anti-tumor immune response (*29*), and specific CD8^+^ T cell states identified by scRNA-seq have been associated with response to immunotherapy in melanoma (*20*). To provide a deeper insight into transcriptional state diversity and cell-state transition of cytotoxic CD8^+^ T cells in ccRCC, we focused on the annotated CD8^+^ clusters (Fig. 2A). Initial clustering analysis of all CD8^+^ T cells (n = 6,529) had identified five distinct clusters of CD8 T cells present across all three patients (Fig.S2A). Examining the cell cycle gene signatures (Fig 2B and S2B) and T cell immune markers of CD8^+^ cell clusters (Fig 2C and S2C; Table S2) revealed three major intra-tumoral cell states: Clusters CD8_0 with increased expression of genes linked to T cell activation and exhaustion (e.g. *CD28, TFRC, TNF* superfamily co-stimulatory members 4, 8 and co-inhibitory receptors *PD-1, TIGIT, LAG3, HAVCR2/TIM-3* and *CTLA4*) compared to others. CD8_3 and CD8_10 displayed increased expression of genes linked to early T cell activation (e.g. *CD44* and *CD69*), cytotoxic (e.g. *IFNG, PRF1, GZMB* and *CCL4*), tissue residency/memory (e.g. *NR4A1, BLIMP1, HOBIT, CCR7* and *TCF7*/*LEF1*) and cell survival (e.g. *FOSB, JUNB, REL, STAT4, STAT1, CD69, CXCR4, IL7R*). Interestingly cluster CD8_17 cells selectively expressed high levels of proliferation marker *MKI67* and were characterized by increased expression of cell cycle genes (*CDK1*/*4* and *CDKN3*). In contrast to intra-tumoral CD8^+^ T cells, peripheral blood cluster CD8_6 cells were enriched for naïve T cell marker *CD62L* (*SELL*) coexisting with cells expressing various levels of cytotoxic markers (e.g. *GZMB, PRF1* and *NKG7*).

**Fig. 2.**
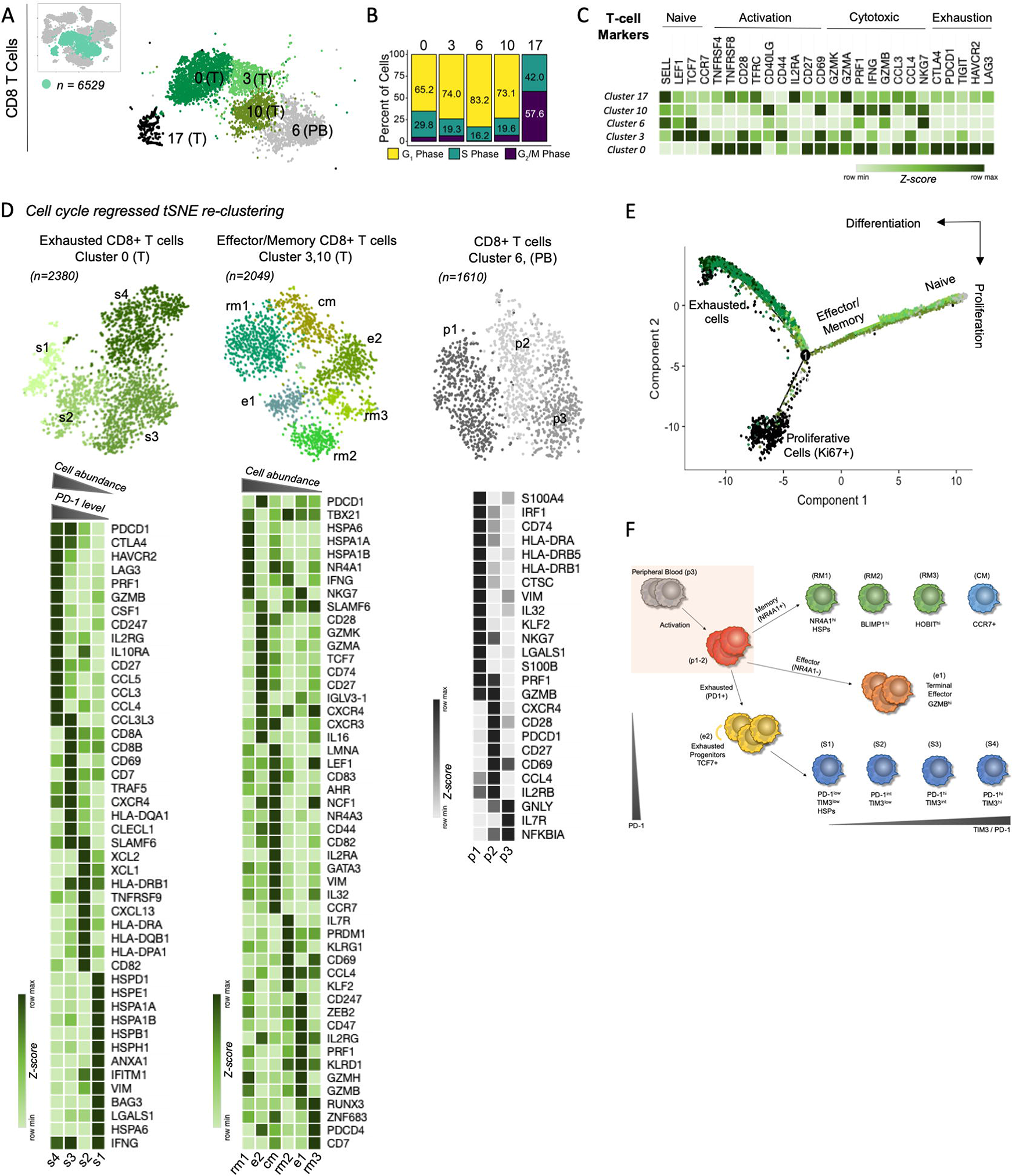
CD8+ T Cells in ccRCC Tumors Exhibit Continuum of Distinct Transcriptional Cell States within Tumor: **(A)** tSNE plot of all CD8+ T cells collected in this study, with cells colored based on five clusters found by *k-* means clustering. **(B)** Assigned cell-cycle state for each cluster. **(C)** Heatmap showing mean expression values of canonical naïve, activated, cytotoxic and exhaustion genes for clusters in (A). **(D)** tSNE plot after cell-cycle regressed tSNE re-clustering of tumor-infiltrating and peripheral blood CD8+ cells found by unsupervised clustering (above). Heatmap showing expression levels for a curated set of transcriptomic signatures identified as differentially expressed and important to exhausted, effector/central memory and activated T cells for CD8 clusters (below). Signature expression are z-scored measured across cell cycle regressed CD8+ T cell clusters. **(E)** Trajectory analysis for the seven CD8+ T cell clusters. Cell expression profiles in a two-dimensional independent space. Each dot represents an individual cell colored by clusters in (A). **(F)** Annotated phenotypic structure for blood and tumor infiltrating CD8+ T cell clusters based on discrete effector, memory and exhausted T cell gene programs.

CD8_0 T cell cluster characterized by the expression of more than one inhibitory receptor e.g. *PDCD1 (*PD-1*), HAVCR2 (*TIM-3*), LAG-3, CD244* or *CTLA4* have been used to describe a state of dysfunction acquired in chronic viral infection and cancer (*30, 31*). These cells represented together more than 50% of the CD45^+^ tumor infiltrating T cells in our ccRCC data (Table S1). To further identify specific cell states and associated gene programs within exhausted CD8_0 T cells, we eliminated effects of cell-cycle and performed dimensional reduction on the cell cycle regressed data. Re-clustering these CD8_0 T cells, we established four distinct subsets of exhausted CD8 T cells each with distinct functional properties (Fig 2D; Table S3). CD8_s1 cells expressed low levels of exhaustion markers *PDCD1 (*PD-1*), CTLA-4, HAVCR2 (*TIM-3*), LAG-3* along with heat shock proteins (*HSPA1A, HSP1B, HSPA6, HSPB1, HSPD1, HSPE1* and *HSPH1).* CD8_s2 cells expressed intermediate levels of *PDCD1* and low levels of *HAVCR2*. In addition, it expressed highest levels of HLA-class II markers. CD8_s3 cells expressed high levels of *PD-1 and CTLA-4* but intermediate levels of TIM-3 while preserving the number of *SLAMF6+* cells, suggesting an early exhaustion-like phenotype. CD8_s4 cells had the phenotype of *PD-1*^hi^ and *TIM-3*^hi^ terminally exhausted sub-population resembling programs in chronic model of viral infection and cancer, with lower proliferation genes (*MKI67, CD8A* and *CD8B)*, high expression of co-stimulatory receptors *4-1BB* and *ICOS* along with cell surface enzyme *ENTPD1 (*CD39), higher cytotoxic markers (*PRF1, GZMB and IFNG)* and increased cytokine production (*CCL3, CCL4, CCL5*, and *CCL3L3)*. Interestingly, using single sample gene set enrichment analysis (ssGSEA), we found Wnt/β-catenin, Notch, Hedgehog signaling, and oxidative phosphorylation pathways enriched in CD8_s2 precursor population of exhausted CD8^+^ T cells (Fig S2D). While exhausted/hsp CD8_s1 sub-population showed pathways enriched for G1S/G2M cell cycle, TGF-β, TNFα via NFκB, IL6-Jak/Stat and Type I (IFNα) and II (IFNγ) interferon signaling indicative of a highly activated state, terminally exhausted CD8_s4 sub-population cells were enriched in both anti- and pro-inflammatory state.

To identify effector/memory-like phenotypes and associated gene programs, we performed dimensional reduction on cell cycle regressed CD8_3 and CD8_10 subsets. Effector and memory (tissue-resident and central) CD8^+^ T cells contribute to promoting effective chronic infection and tumor immunity (*32, 33*). Examining the genes associated with memory precursors (*KLRG1* and *IL7R*), genes involved in the regulation of resident memory (*NR4A1, ZNF683*/*HOBIT, PRDM1*/*BLIMP1, RUNX3* and *CD69*) and central memory (*CCR7* and *SELL*), cytotoxic effector molecules granzymes A, B, K and H (*GZMA, GZMB, GZMK* and *GZMH)* and cytokine receptor subunits (*IL2RG)*, we annotated six effector/memory clusters (Fig 2D Table S4). Our data shows that effector and memory clusters were distinguished by the expression of nuclear family receptor for tissue residence, *NR4A1.* Variable levels of expression of *NR4A1* gene were observed in resident-memory CD8 clusters, highest in cells expressing heat shock protein (CD8_rm1), intermediate in (CD8_rm2) cells expressing high levels of *BLIMP1* and lowest in (CD8_rm3) cells expressing high levels of *HOBIT and RUNX3*. Interestingly, CD8_rm2 cells had increased expression of *IL7R, KLRG1, CD69* and *KLF2* genes associated with resident-memory differentiation and maintenance program (*34, 35*) compared to others. CD8_cm cells were enriched in *CCR7* along with increased expression levels of activation and proinflammatory molecules (*CD44, IL32* and *IL2RA*). Effector cells expressing granzymes and perforins, split into two sub-clusters, CD8_e1 (*GZMB, GZMH* and *PRF)* and CD8_e2 (*GZMA* and *GZMK).* We labelled CD8_e1 cluster as cell undergoing terminal differentiation that expressed high levels of perforins and granzymes with effector genes *KLRG1* and *PDCD1* but do not express surface markers *SELL*, *CCR7*, *CD27* and *CD28* as described in (*29*). CD8_e2 appeared to have enrichment of *TCF7*, a transcription factor associated with expansion of progenitor exhausted CD8^+^ T cells wherein an increased ratio of progenitor to terminally exhausted cells is correlated with responsiveness to PD1-therapy in melanoma patients (*20*). Similarly, *TCF1* (mouse homolog of h*TCF7*) is also known as a driver of pre-exhausted CD8 progenitor population in chronic viral infection (*36*). The CD8_e2 subset also expressed intermediate levels of *PDCD1* and higher *SLAMF6*, consistent with the phenotype of progenitor population of exhausted T cells as described in the literature (*37, 38*). In addition, our ssGSEA analysis found that effector and central/resident memory sub-clusters exhibit different levels of genes contributing to signatures for CD8 activation inflammatory-pathway related genes for IL2/Stat5, IL6-Jak/Stat, TNFα via NFκB, Type II (IFNγ) interferon signaling and anti-inflammatory signaling (Fig S2E). As reported in (*39*), the central memory sub-population (cm) showed enrichment in canonical Wnt/β-catenin pathway along with Type I (IFNγ) interferon signaling. Interestingly, the progenitor exhausted (e2) and the resident memory sub-population (rm3), are enriched for gene signatures associated with TCA cycle, oxidative phosphorylation and lipid mediators as their major metabolism pathway (Fig S2E), as previously described (*40*).

To identify gene programs distinguishing the peripheral blood of ccRCC patients, we examined cluster (CD8_6) post cell cycle regression. We defined three distinct clusters (CD8_p1, CD8_p2, CD8_p3) (Fig 2D; Table S5). Interestingly, *GZMB* and *PRF1* had much higher expression in CD8_p1 and CD8_p2 with the latter expressing higher levels of co-inhibitory receptor *PDCD1*, co-stimulatory molecules (*CD27* and *CD28)*, cytokine (CCL4), cytokine receptor (*IL2RB)* and chemokine receptor *(CXCR4).*We next performed pseudotime analysis ordering cells by trajectory of gene expression to define the transitions between cell states and potential branch points between peripheral blood and intratumor CD8^+^ T cells (*41*). CD8^+^ T cells were distributed along the pseudo-temporally ordered paths from naïve CD8^+^ T cells in peripheral blood to effector/memory cells infiltrating-tumor and branching into exhausted or terminally differentiated cells reflecting a possible path for differentiation. Another side branch led into *MKI67*^+^ CD8^+^ cells suggesting proliferative cell state arises from effector/memory CD8 T cells upon antigen stimulation (Fig 2E and S2F).

### In-depth Characterization of CD4 T Cells in ccRCC Tumors Reveals Effector, Memory and Regulatory T Cell Sub-phenotypes

CD4^+^ T cells can target the tumor cells in various ways, either directly through cytolytic mechanisms or indirectly by modulating the tumor immune microenvironment. While CD4^+^ T cells promote T cell priming as well as the effector and memory functions of CD8^+^ T cells, regulatory CD4^+^ T (Tregs) cells play key role for dampening responses from immune system against cancer. Transcriptome profiling from our matched tumor and blood data revealed five clusters of CD4^+^ T cells present across all three patients (Fig 3A and S3A; Table S6). Intratumoral CD4_21 cells selectively expressed high levels of cytolytic gene programs including granzymes (*GZMA, GZMB, GZMK* and *GZMH*), pro-inflammatory cytokines (*IFNG and IL32*), chemokines (*CCL4* and *CCL5)*, cytokine/chemokine receptors (*IL2RB, IFNGR1, CXCR3* and *CXCR4)* and exhaustion markers *(PDCD1* and *LAG3)* (Fig 3C and S3C). CD4_1, CD4_7 and CD4_12 cells in the peripheral blood were distinguished by an exclusive expression of naïve T cell markers (*SELL, TCF7*/*LEF1* and *CCR7*). A single cluster of *FOXP3*^+^ Tregs, *i.e.* CD4_14, was identified, which contained both tumor-infiltrating and peripheral blood Treg subpopulations. On visualizing the cell cycle state, we observed that Tregs (CD4_14) (circulating and intra-tumoral) and intra-tumoral CD4^+^ T cells (CD4_21) exhibited elevated S/G2M percentages (indicated by green and purple, respectively), indicative of an active state of proliferation (Fig 3B and S3B).

**Fig. 3.**
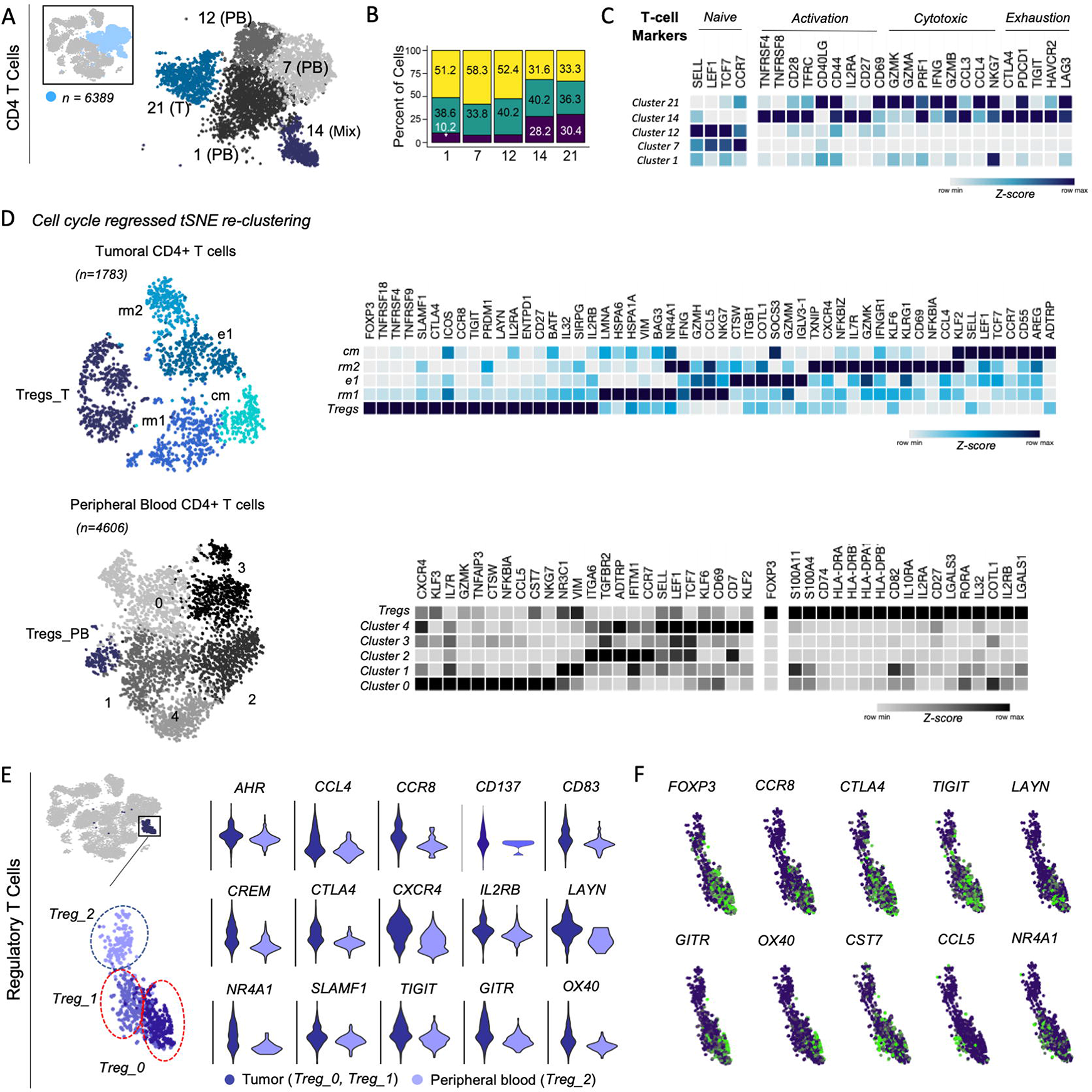
In-depth Characterization of CD4 T cells in ccRCC Tumors Reveals Effector, Memory and Regulatory T Cell Sub-phenotypes: **(A)** tSNE plot of all CD4+ T cells collected in this study, with cells colored based on five clusters found by *k-* means clustering. **(B)** Assigned cell-cycle state for each cluster. **(C)** Heatmap showing mean expression values of canonical naïve, activated, cytotoxic and exhaustion genes for clusters in (A). (D) tSNE plot after cell-cycle regressed tSNE re-clustering of both, tumor-infiltrating and peripheral blood CD4+ cells. **(D)** tSNE plot after cell-cycle regressed tSNE re-clustering of tumor-infiltrating and peripheral blood CD4+ cells found by unsupervised clustering (left). Heatmap showing expression levels for a curated set of transcriptomic signatures identified as differentially expressed and important to effector, memory and regulatory T cells for CD4 clusters (right). Signature expression are z-scores measured across cell cycle regressed CD4+ T cell clusters. **(E)** tSNE plot shows Treg subsets (left) and comparison of differential genes expressed in tumor infiltrating Tregs versus peripheral blood (right). **(F)** Key genes enriched between tumor-infiltrating two Treg subsets (Treg_0 and Treg_1) as defined in (Table S10).

To analyze cellular identity and differentiation programs within intratumoral cytotoxic (CD4_21) and Tregs cells (CD4_14, tumor subset), we analyzed the substructure of these cells after cell cycle regression. Re-clustering analysis revealed total of five intra-tumoral transcriptional states for CD4^+^ cells (Fig 3D; Table S7). We evaluated distinguished gene components across these discrete clusters to identify memory and effector gene programs. Our data showed two resident memory cluster; CD4_rm1 cells highly expressed resident memory markers (*NR4A1*, *HOBIT, RUNX3)* and proinflammatory molecules (*GZMH* and *IFNG)* along with heat shock proteins (*HSPA1A, HSPA1B and HSPA6)*; whereas CD4_rm2 cells expressed higher transcript levels of *CD69, IL7R, KLRG1, KLF2, KLF6* and granzyme *GZMK*. CD4_cm cell markers were consistent with central memory-like phenotype (*CCR7* and *TCF7*/*LEF1*) enriched in genes for Wnt/β-catenin signaling pathway, while CD4_e1 had the phenotypes of early effector cells (*IGLV3-1, CTSW, COTL1* and *ITGB1*) identified during CD8+ T cell analysis. Re-clustering analysis of cell cycle regressed CD4^+^ T cells from peripheral blood revealed a total of six states with distinct naïve (clusters CD4_1, CD4_2, CD4_3 and CD4_4), effector (cluster CD4_0) and regulatory (cluster Tregs_PB) phenotype (Fig 3D; Table S8). In addition, we used trajectory analysis to identify a transition of peripheral blood CD4+ T cells into cytotoxic tumor-infiltrating cells and identified a main trajectory branch arising from peripheral blood and giving rise to effector/memory cell states in the tumor.

Selectively targeting immunosuppressive cancer associated Tregs while maintaining peripheral immunity has been of considerable interest following FDA approval of anti-CTLA4 mAb in cancer therapeutics. Tregs in the tumor exclusively expressed *FOXP3*, with high expression of both co-inhibitory immune checkpoints (*CTLA-4 and TIGIT)*, and co-stimulatory molecules *(CD27, CD28, ICOS, GITR* and *OX40)* (Fig 3D). In order to identify distinct immune checkpoint markers expressed on Tregs subset in the ccRCC tumor microenvironment, we compared transcriptomic expression of tumor infiltrating Tregs versus peripheral blood. scRNA-seq analysis revealed that Treg_T distinguished from Treg_PB by its increased expression of *CCR8, NR4A1, CTLA4, ICOS* and higher expression of *GITR, OX40, CD137, TIGIT, BATF, IL32* and *HLA* molecules (Table S9). Interestingly, Tregs with tumor revealed two Treg states (Treg_0 and Treg_1) (Fig 3F; Table S10). Treg_0 cells expressed higher transcripts for the Treg master regulators *FOXP3, CCR8, CTLA4, TIGIT* along with co-stimulatory molecules GITR and OX40, whereas Treg_1 Tregs expressed *CCL5* (*RANTES*) and *NR4A3*. The gene expression program of Treg_0 strongly corresponds to superior immunosuppressive state of these Tregs compared to their blood and tumor counterparts. Comparing intratumoral Tregs and cytotoxic CD4^+^ T cells; in addition to increased anti-inflammatory gene program, the Tregs displayed an enrichment in metabolic pathway genes set associated with glycolysis, glycogen metabolism and TCA cycle to fuel oxidative phosphorylation (Fig. S3E) required for Treg suppressive function (*42*).

### Enriched TCR sequences in Subsets of T cells from Paired Tumor and Peripheral Blood

We reconstructed the T-cell receptor (TCR) sequences (α and β chains) for CD8^+^ and CD4^+^ T cells to evaluate TCR clonality based on shared CDR3 sequences in both α and β chains, as previously described (*20*). We first analyzed the abundance of each TCR clonotype observed in each patient (blood and tumor) (Fig. S4). After filtering to exclude sequence errors, the estimate total number of T cells with productive combined TCRα and TCRβ was lower compared to the overall T cells in each tissue. The abundance of productive clonotypes is generally greater in blood, but the cumulative abundance within the top 10 clonotypes is higher in tumor for each patient (Fig. S4). These metrics indicate that tumors have less diverse clonotypes than those in the blood. To systematically assess the intratumoral TCR clonotypes and CDR3 sequence overlap with peripheral blood, we defined the fraction of T cells in tumor sample with at least one identical clonotype in the peripheral blood and calculated over all three pairs of samples. Calculating the interpatient tissue combinations at the sequencing depth used in this study, approximately less than 5% clonotypes in tumor and blood overlap with each other. Despite the significant fraction of overlapping T cell clonotypes between tumor and blood within the same patient, some highly abundant clonotypes in the tumor were either not detected or detected at very low levels in blood (and vice versa) (Fig. S4). No identical TCR sequences were found between tissues from different patients (Fig. 4A). However, some degree of CDR3 sequence overlap between tumor and blood from different patients (Fig. 4B). The extent of the overlap is variable from patient to patient, but the trends are similar. Since peripheral blood is more accessible tissue and the tumor-clonotypes present in blood could be useful for future diagnostic or therapeutic purposes, we reveal the top five shared clonotypes in the tumor and peripheral blood of each patients along with CDR3 amino acid sequence (Fig. 4C). Next, we analyzed the enrichment of TCR clonotype and more than one TCRs were detected in more than one T cell in a single sample (n=2; doublets and n=3; triplets) compared to single TCRs found in one T cell of each patient. Interestingly, enriched TCRs were more common in intratumoral CD8^+^ T cells compared to single TCRs in peripheral blood (Fig. S5). This was observed for both CD4 and CD8^+^ T cells suggesting that the tumor T cells were likely exposed to repeated tumor antigen stimulation leading to clonal expansion of certain dominant clones. Lastly, to understand the T cell dynamics and TCR plasticity between transcriptional subsets of T cells identified, we revealed an overlap of TCR clonotypes within original CD8 and CD4 cluster groups (Fig. 4D). We identified a substantial overlap of T cell receptors among different subsets of CD8^+^ T cells within both tumor and peripheral blood. In contrast, we only observed significant number of shared clonotypes within the tumor-infiltrating Tregs and cytotoxic CD4^+^ T cells. Interestingly, Tregs demonstrated the highest proportion of shared TCR diversity compared with other T cell states. While averaged 36.6% clonotypes among all three patients were Treg specific, the Treg cluster shared 32.6% clonotypes with effector T cell clusters and 30.8% with naïve population clusters.

**Fig. 4.**
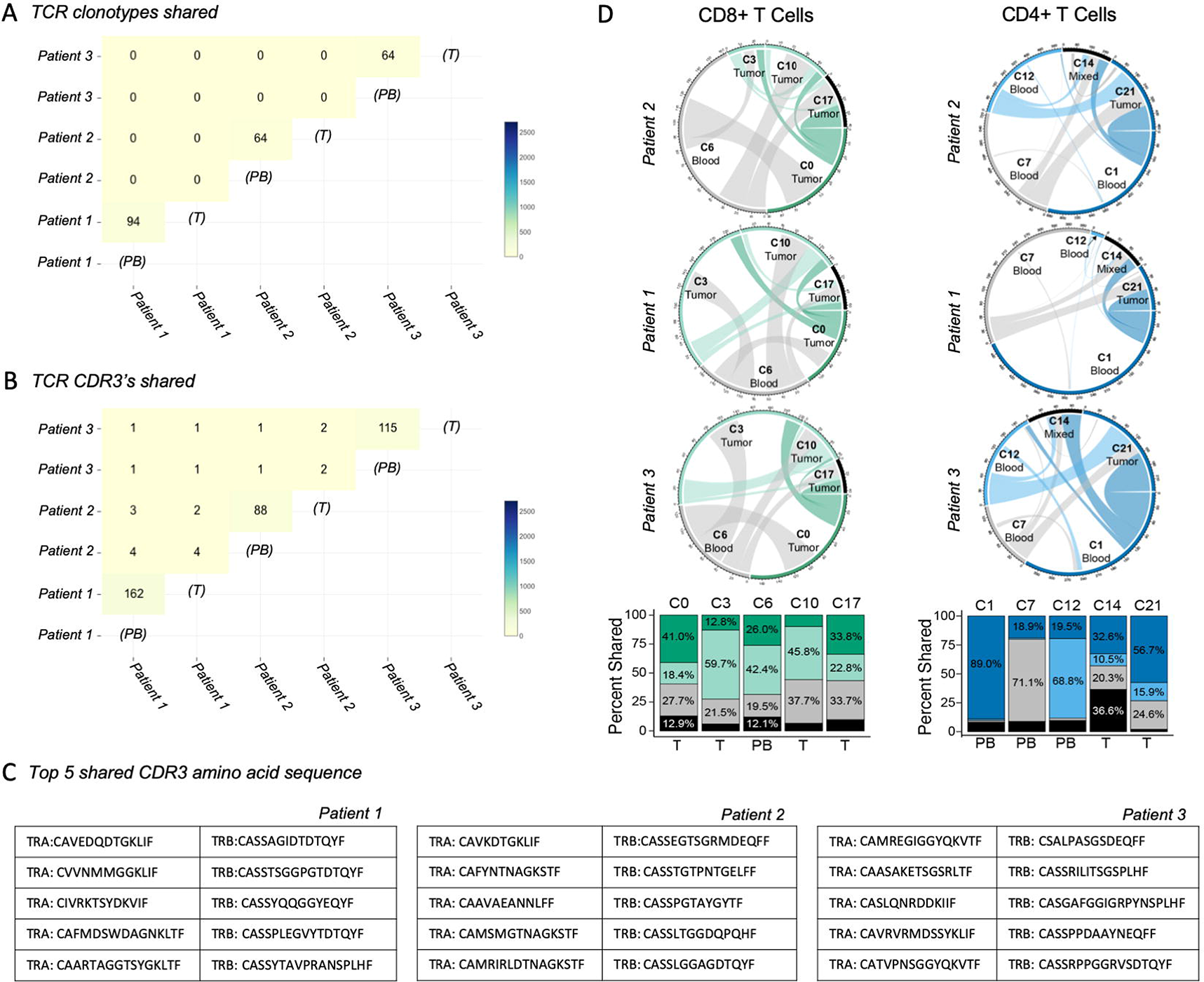
Enriched TCR Sequences in Patient Blood Partially Overlap with Those in their Tumors and are Shared between Subsets of T cells: **(A)** TCR clonotypes shared between ccRCC tumor and blood tissue across all three-patients (Morisita-Horn metric). **(B)** CDR3s shared across blood and tumor of all three-patients (Morisita-Horn metric). **(C)** CDR3s of top 5 shared clonotypes of individual patients. **(D)** TCR receptor shared between TCR receptor shared between clusters of CD8 and CD4 T cells by patient (chord diagrams) and mean TCR receptors shared between clusters across all three patients (bar chart).

### Single-cell Transcriptional Analysis of Tumor-associated Macrophages and Dendritic Cells Reveals Distinct Subsets with Unique Marker Genes

Using the same clustering analysis, ccRCC myeloid cells expressing *CD68* gene, partitioned into five major cell populations across tumor and blood, present in all three patients (Fig 5A; Table S11). Signature of genes of monocytes (*CD14 and CD16)* were enriched in cluster 5, 9 and 16. Cluster 4 cells were enriched with tumor-associated macrophage signature genes such as *APOE*, *APOC1*, *MRC1*; complement genes (*C1QA*, *C1QB* and *C1QC*); and cathepsin *CTSB*; and cluster 13 with dendritic cell specific genes *CD1c* and *FCER1A.* Monocytes present within both tumor and peripheral blood can be split into two subsets, comprised of *CD14*^+^ *CD16*^−^ ‘classical’ monocytes (mono1 and mono3) and *CD14*^−^ *CD16*^+^ ‘non-classical’ monocytes (mono2 and mono3) (*43*) (Fig 5B and S6B). To investigate if tumor microenvironment altered transcriptional phenotype of these monocytes, we performed differential gene expression and ssGSEA pathway analysis (Fig S6C; Table S12). Both tumoral-associated monocytes, mono1 and mono2 were enriched in inflammatory cytokine and chemokines (*IL1b, CCL4, CCL3L2* and *CCL4L2)* along with heat shock proteins (HSPs). In addition, the classical monocytes within tumor discretely expressed *TREM1, EGR1, AREG, MAP3K8, NLRP3*, chemokines (*CCL2, CXCL2* and *CXCL8)* and suppressor of cytokine signaling (*SOCS3)*; non-classical intratumoral monocytes were enriched in anti-apoptotic molecules (*RIPK2, BCL2 and BIRC3*), tissue resident markers (*NR4A1* and *NR4A2*), NFκB signaling genes (*NFKBIA* and *NFKBIZ)*, chemokines (*CXCL16* and *CCL4)* and receptor (*CXCR4*) (Fig 5B).

**Fig. 5.**
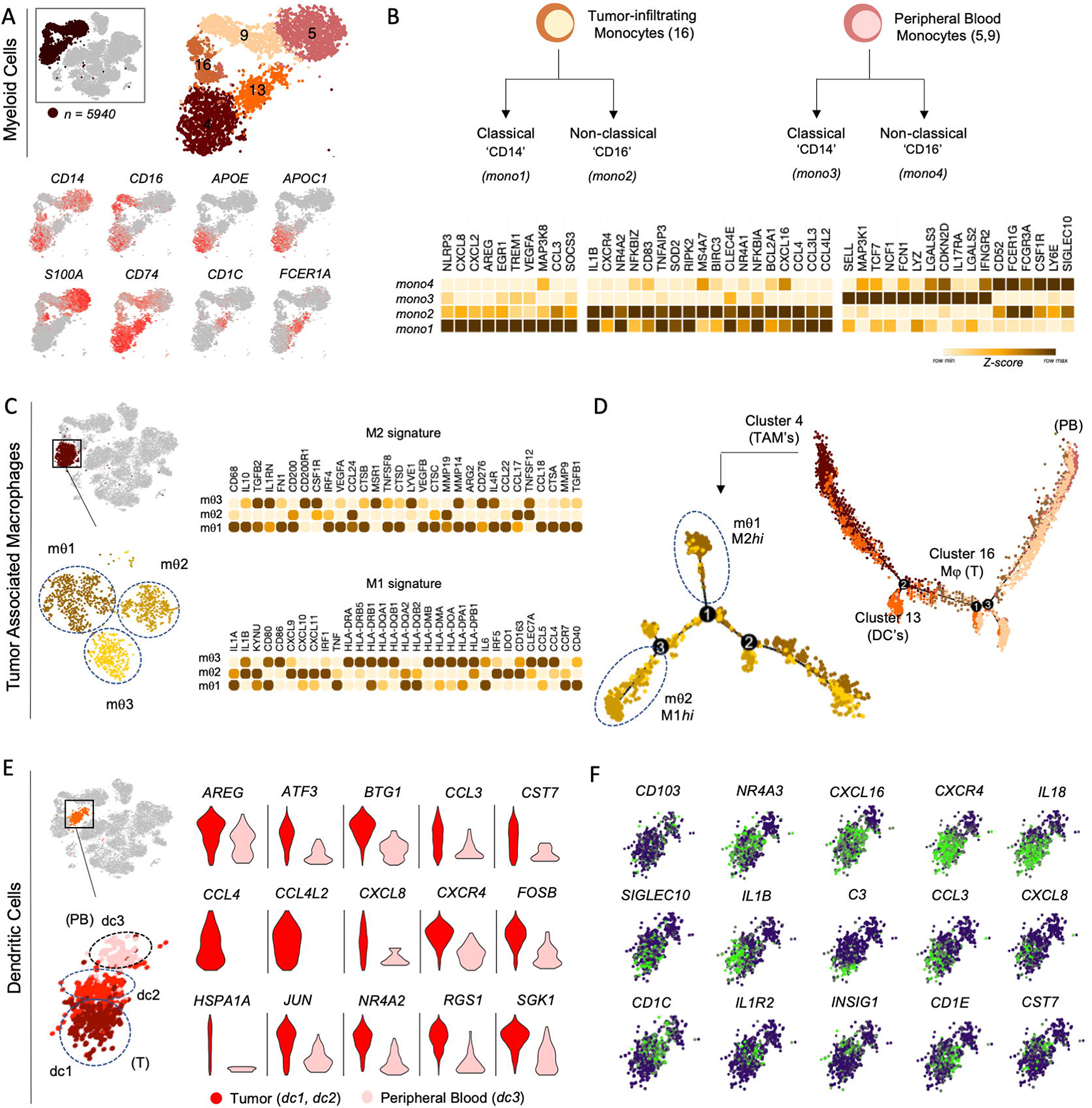
Single-Cell Transcriptional Analysis of Tumor-associate Macrophages and Dendritic Cells Reveals Distinct Subsets with Unique Marker Genes: **(A**) tSNE plot of only myeloid cells across all-three patients. Cells are colored based on five clusters found by unsupervised clustering (above). tSNE plot showing expression levels for a curated set of transcriptomic signatures important to myeloid cells (below). **(B)** Annotated phenotypic structure for ‘classical’ and ‘non-classical’ monocytes within tumor and in peripheral blood. Heatmap showing z-scored mean expression values of discriminating genes for identified four monocytes clusters. **(C)** tSNE plot shows TAM subsets (left) and heatmap showing scaled expression values of discriminating M1/M2 signature genes expressed for TAM clusters (right). **(D)** Trajectory analysis for the five myeloid cell clusters. **(E)** tSNE plot shows DC subsets (left) and comparison of differential genes expressed in tumor infiltrating Tregs versus peripheral blood (right). **(F)** Key genes important to dendritic cells and enriched between tumor-infiltrating two DC subsets (dc1 and dc2) as defined in (Table S15).

We defined a total of 3 transcriptional states of tumor associated macrophages corresponding to cluster_4 (mθ1-3) (Fig 5C and S6E; Table S13). We examined gene signatures to assess whether any of the states corresponds to the defined M1 and M2 states, which have been used to define pro-inflammatory and anti-inflammatory macrophages in vitro, respectively. mθ1 macrophages, the largest macrophage population exhibited enrichment for M2 signature (*CD68, CD200, ARG2, IL4R, TGFB1, TGFB2, IL10* and *FN1)* while mθ2 showed an increase in the M1 gene signature (*IL1B, KYNU, CD80, CD86, IDO1, CD163* and *HLA* antigen presentation genes). However, mθ3 cells expressed both, suggesting that while M1 and M2 are distinct cell states in ccRCC tumors, the macrophage polarization is a dynamic process. This phenomenon is also well represented by the pseudo-time plot of tumor-associated monocyte/macrophages (Fig. 5D and S6F).

Clustering of DCs (Cluster 13) identified two subsets of tumor DCs (dc1 and dc2) and one peripheral blood (dc3) with distinct gene expression programs in each (Fig. 5E). The DC subsets were present in all three patients despite the low cell numbers in this cluster. To assess intratumoral DCs, we compared their gene expression program with peripheral blood DCs (Fig. 5E Table S14). The tumoral DCs uniquely overexpressed immune cell maturation and proliferation *CSFR*, antigen presentation *HLA* molecules, immune modulators including *CD83, CD86* and *TIM3* and cell cycling molecule *CDKN1A.* The tumoral DCs also expressed high amounts of cell adhesion molecules (*CD9* and *CD81)*, cytokines (*1L18* and *IL1B)*; chemokine receptor (*CXCR4)*, and Fc and complement receptors (*CD32* and *CD11b)*. Both tumor and blood DCs contained few cells expressing *ITGAE* (*CD103*). To assess tumoral DC heterogeneity, we compared gene expression programs across the three DC subsets (Fig. 5F, Table S15). The dc1 subset expressed activated DC phenotype uniquely expressing *CD69, CCL3, CCL4, CXCL2, CXCL8, IL18, IL1B, SIGLEC10 and CLEC9A* as well as complement receptors (*C3, C1QA, C1QB* and *C1QC)*. The dc2 cells highly expressed canonical DC markers like *CD1C, CD1E* in addition to immune associated genes (*IL1R2, CST7*, *INSIG* and *CLEC4C).*

## Discussion

Given the success of ICB in ccRCC, there has been renewed interest in understanding the features of the tumor immune microenvironment that promote response (or resistance) to cancer immunotherapy (*44*). Improved understanding of the immune contexture of ccRCC will facilitate investigation into the roles and regulation of discrete immune cell states governing response and resistance to cancer therapies. To provide a comprehensive analysis of the immune landscape of ccRCC, we performed single cell RNA-sequencing of tumors and peripheral blood from three patients with treatment-naïve, localized ccRCC.

CD8^+^ T cells are key effectors of anti-tumor immunity (*29*). PD-1 blockade disrupts adaptive immune resistance resulting in increased intratumor infiltration of CD8^+^ T cells (*45*). Recently, scRNA-seq profiling of melanoma has revealed unique CD8 T cell states associated with response and resistance to PD-1 blockade (*20*). Unlike melanoma and most other cancers, CD8^+^ T cell infiltration portends a worse prognosis in ccRCC (*12, 46*), suggesting the balance of CD8^+^ T cell states may be altered in ccRCC compared to other cancer types. Here, we identified multiple intratumoral CD8 T cell states that exhibit hallmarks of effector/memory (CD8_3 and CD8_10) and exhausted (CD8_0) subpopulations that are generated after antigenic stimulation. As recently identified in melanoma (*20*), surface expression of *ENTPD1* (CD39) and *TIM3* readily discriminated between these two CD8 sub-populations, with CD39^−^ TIM3^−^ CD8 T cells enriched for memory/effector-like genes, while CD39^+^ TIM3^+^ CD8 T cells represented a terminally exhausted state. Interestingly, these subtypes formed a smooth continuum with activated CD8 T cell states observed in peripheral blood. We demonstrated that the relative abundance of exhausted CD8^+^ T cell infiltrates in ccRCC (defined by *PD-1, TIM-3*) was highest out of all observed distinct cell states and observed in all three patients. These cells were not observed in peripheral blood. The high prevalence of exhausted CD8^+^ T cell infiltrates may partially explain the sensitivity of ccRCC to ICBs. Interestingly, we detected a high frequency of *HLA-DR* expressing T cells within the *PD-1*^+^ *TIM3*^+^ enriched CD8 T cell clusters, which have been previously identified as CD8^+^ HLA-DR^+^ regulatory T cells (*47*). Within the effector/memory subset, progenitor exhausted CD8^+^ T cells (expressing *TCF7* and *SLAMF6*) represented a small proportion of the total evaluated CD8 T cells in our data. As *TCF7*^+^ *PD-1*^+^ CD8^+^ T cells have been associated with response to PD-1 blockade in melanoma (*20*), it is tempting to speculate that the paucity of *TCF7*^+^ (e2) CD8^+^ T cells relative to early and terminal exhausted (s3, s4) contributes to the poorer prognosis observed in patient with heavily CD8^+^ T cell infiltrated ccRCC. Although the functional role of *TCF7*^+^ CD8 subset in ccRCC is unclear, intratumoral expansion of this progenitor exhausted CD8 population has been reported to derive from T cell clones that may have just entered the tumor after ICB therapy in immune-responsive cancers (*48*). The recent independent CyTOF study that utilized single-cell multiplexed profiling of CD8^+^ T cells from cohort of 73 ccRCC patients (*16*) was insufficient in capturing full phenotypic heterogeneity, thus highlighting the power of single-cell transcriptome mapping. Future studies will be required to evaluate the contribution of the observed CD8^+^ T cells to response or resistance to PD-1 blockade in ccRCC, as well as how the CD8^+^ T cell compartment evolves over time following treatment with immune checkpoint blockade +/− anti-angiogenic therapy.

The pro- and anti-tumor role of CD4^+^ T cells still remains unexplored and unappreciated in the TME of ccRCC. Here, we demonstrate intratumoral Foxp3-cytotoxic and Foxp3+ regulatory CD4 T cell clusters, define transcripts distinguishing them from their peripheral blood counterpart and establish tumor-infiltrating effector and memory programs in addition to naïve cell states in peripheral blood. Surprisingly, very few of the effector or memory CD4^+^ T cells expressed *PD-1* gene (Fig. S3), suggesting impaired antigen recognition and/or priming. Moreover, circulating CD4^+^ TCR repertoire showed less diversity compared with circulating CD8^+^. This is of particular interest given that greater diversity of CD4^+^ blood T-cell clones before immunotherapy has been significantly correlated with long-term survival upon CTLA4 or PD-1 inhibition in melanoma (*49*). We revealed that a substantial portion of intratumoral Tregs showed higher expression of *FOXP3* and other transcripts for effector T cell suppression genes involved in essential Treg function, as recently reported in a non-tumor context (*50*). The single cell data provided us with an opportunity to correlate TCR sequence abundance and clonal diversity with the transcriptional output of each T cell cluster. Looking at the TCR clonotype in tumor and blood, we discovered enriched CDR3 amino acid sequence in individual tissue along with shared sequences that could aid in investigating ccRCC specific tumor neo-antigens.

Clonotype analysis showed a significant overlap in TCRs between Tregs and non-Treg CD4^+^ T cells, indicative of a potential conversion of non-Treg CD4^+^ T cell to Tregs, as proposed by other groups (*51, 52*). However, the majority of Tregs have distinct TCRs that do not overlap with non-Tregs, which implies their thymic origin and could be stimulated by tumor-originated self-antigens for clonal expansion. Also, it suggests that Tregs can suppress effector T cells in both antigen-specific and antigen-independent fashion. Shared clonality may provide superior antigen-specific suppression of effector T cells by Tregs, which is further strengthened by antigen-independent suppression to collectively provide an immune suppressive TME.

High numbers of immunosuppressive myeloid derived cells in the peripheral blood and tumors have been shown to associate with a poor clinical outcome in ccRCC (*53*). We observed a heterogeneous myeloid landscape in tumor and peripheral blood of ccRCC patients. We demonstrated that both classical and non-classical subsets of peripheral blood monocytes exist within the ccRCC TME, however at low frequency and expressed distinguished gene programs. We also observed multiple cell states for both TAMs and dendritic cell types. The observed cell states were shared across all three patients, and each cell sub-type demonstrated gene programs that form a continuum. We examined distinct gene signatures to assess whether any of the TAM states correspond to canonical M1 and M2 states which have been used to define anti-inflammatory and pro-inflammatory macrophages (*54*). While the TAM clusters exhibited a dominant M1-like or M2-like gene programs, the presence of an intermediate cluster (mθ3) showing shared features of M1 and M2 represents the transcriptional macrophage plasticity. In contrast to the identification of 17 TAM phenotypes, our pseudotime evaluation of the myeloid compartment by scRNA-seq discovered six distinct cell states. Future studies will be required to confirm and extend these findings. Lastly, our data identified the gene expression programs of monocyte-derived DCs in ccRCC tumor, most of which corresponded to the hDC subsets recently reported in lung cancer (*55*).

This study highlights the complexity and diversity of ccRCC TME predominantly infiltrated by CD8^+^ T cell and TAMs. What emerges from the focused evaluation of the diversity of lymphoid and myeloid cell states is a preponderance of immune suppressive cell populations/states. The relative abundance of exhausted/dysfunctional CD8 cell state, CD4^+^ Tregs, and M2-like TAMs is indicative of a highly immunosuppressive TME. While our study represents the first report characterizing the immune landscape of ccRCC using sc-RNAseq, it has several limitations. First, despite the large number of single cells characterized, tumors and matched blood were analyzed from only three patients. It is notable that though the three patients showed variable proportions of lymphoid and myeloid cells in each cell state, the number of states remain consistent across for each patient. Evaluation of additional patients will be needed to determine if the observed cell states are representative of a wider population and would profiling more patients reveal additional immune cell subsets. It is possible that rare immune cell subsets were not captured, although even rare subsets like B lymphocytic plasma cells (representing less than 1% of all cells) were detected in all three patients (Fig. 1A-B), suggesting that rare cell populations could readily be identified and deeply characterized in addition to the broader lymphocyte and myeloid transcriptional cell states. Nonetheless, analyzing more patients could refine immune cell states observed here. Secondly, the immune landscape described is exclusively from treatment-naïve ccRCC patients. Follow up studies will be needed to identify cell states associated with response and/or resistance to immune checkpoint blockade +/− anti-angiogenic therapy. Furthermore, any novel findings will require validation using tissue imaging in larger and independent cohorts to correlate the abundance of discovered cell phenotype to clinical outcomes and therapeutic responses to immunotherapy. For example, in the study of CD8 T cell states associated with response to PD-1 blockade, Sade-Feldman and colleagues validated *TCF7* protein expression in CD8 T cells by multiplexed immunofluorescence (*20*). Thirdly, functional profiling of specific cell states in a tumor model will be needed to define the roles and regulation of the identified lymphoid and myeloid sub-populations to tumor initiation, progression, metastasis, and response/resistance to anti-angiogenic and immune therapies in ccRCC (*20, 56*). Nevertheless, the immune landscape of ccRCC reported here will provide a valuable resource to accelerate research of the TME with the ultimate goal of discovering novel transcriptomic biomarkers and therapeutic targets.

## Materials and Methods

### EXPERIMENTAL MODEL

#### Subject Details and Tissue Collection

Fresh blood and primary clear cell renal cell carcinoma (ccRCC)samples were obtained from the University of Iowa Tissue Procurement Core and GUMER repository through the Holden Comprehensive Cancer Center from subjects providing written consent approved by the University of Iowa ethics board committee. The patients ranged from 67 to 74 years old; the tumor samples were of diverse tumor stages and sourced from male subjects. Tumor grades were histologically determined by a pathologist. Three ccRCC tumor specimens paired with individual blood samples were used in the study.

#### Tumor Dissociation and Isolation of Mononuclear Cells

Renal tumor samples were dissociated into single cells by a semi-automated combined mechanical/enzymatic process. The tumor tissue was cut into pieces of (2-3mm) in size and transferred to C Tubes (Miltenyi Biotech) containing a mix of Enzymes H, R and A (Tumor Dissociation Kit, human; Miltenyi Biotech). Mechanical dissociation was accomplished by performing three consecutive automated steps on the gentleMACS Dissociator (h_tumor_01, h_tumor_02 and h_tumor_03). To allow for enzymatic digestion, the C tube was rotated continuously for 30 min at 37°C, after the first and second mechanical dissociation step (*57*). Cells from fresh tumor specimens were incubated with FcR blocking reagent (StemCell Technologies) for 10 min at 4°C and labelled with 1ug/ml of the FITC anti-human CD45 antibody (BioLegend) per 10^7^ cells for 20 min at 4°C. CD45+ cells were isolated using the EasySep™ FITC Positive Selection Kit (StemCell Technologies). Alternatively, mononuclear cells (MNCs) from whole peripheral blood of paired subjects were isolated using SepMate Tubes (StemCell Technologies) by density gradient centrifugation. Cells were then viably frozen in 5% DMSO in RPMI complemented with 95% FBS. Cryopreserved cells were resuscitated for flow cytometry analyses by rapid thawing and slow dilution.

#### Cell Sorting for Single Cell RNA-sequencing

Viable immune (CD45+ Hoechst-) single cell suspensions generated from three ccRCC tumor samples and blood were FACS sorted on a FACS ARIA sorter (BD Biosciences) for lymphoid and myeloid cells (Ratio 3:1). The cells were sorted into ice cold Dulbecco’s PBS + 0.04% non-acetylated BSA (New England BioLabs). Sorted cells were then counted and assessed viability MoxiGoII counter (Orflo Technologies) ensuring that cells were re-suspended at 1000cells/ul with a viability >90%.

#### RNA-seq 10X Genomics Library Preparation, Single Cell 5’ and TCR Sequencing

Single-cell library preparation was carried out as per the 10X Genomics Chromium Single Cell 5’ Library and Gel Bead Kit v2 #1000014 (10 × Genomics, Pleasanton, CA, USA). Cell suspensions were loaded onto a Chromium Single-Cell Chip along with the reverse transcription (RT) master mix and single cell 5′ gel beads, aiming for 7,500 cells per channel. Following generation of single-cell gel bead-in-emulsions (GEMs), reverse transcription was performed using a C1000 Touch Thermal Cycler (Bio-Rad Laboratories, Hercules, CA, USA). 13 cycles used for cDNA amplification. Amplified cDNA was purified using SPRI select beads (Beckman Coulter, Lane Cove, NSW, Australia) as per the manufacturer’s recommended parameters. Post-cDNA amplification reaction QC and quantification was performed on the Agilent 2100 Bioanalyzer using the DNA High Sensitivity chip. For input into the gene expression library construction, 50ng cDNA and 14 cycles was used. To obtain TCR repertoire profile, VDJ enrichment was carried out as per the Chromium Single Cell V(D)J Enrichment Kit, Human T Cell #1000005 (10 × Genomics, Pleasanton, CA, USA) using the same input sample. Sequencing libraries were generated with unique sample indices (SI) for each sample and quantified. Libraries were sequenced on an Illumina HiSeq 4000 using a 150 Paired End sequencing kit.

### QUANTIFICATION AND STATISTICAL ANALYSIS

#### Pre-processing of Paired 5’ Single-cell RNA-seq Data and Initial Analysis

Basecalls were converted into FASTQs using the Illumina bcl2fastq software by the University of Iowa Genomics Division. FASTQ files were aligned to human genome (GRCh38) using the Cell Ranger software pipeline (version 2.2) provided by 10xGenomics using the STAR aligner as described by manufacturer (*58*). Filtering read alignments and cellular barcode and Unique Molecular Identifier (UMI) counting to determine gene-barcode matrices for each sample were also performed by Cell Ranger. Reads from tumor and blood sample of individual patients were aggregated and libraries were normalized to the same sequencing depth using *cellranger aggr* function to generate normalized aggregate data for each subject. Secondary clustering and gene expression analysis was performed by the Loupe Cell browser. Secondary analysis of gene expression (*i.e.*, dimensionality reduction, graph-based clustering, K-means clustering and gene expression analysis specific to each cluster) was also performed by the Cell Ranger software using default settings for a rough secondary analysis. Initial data exploration was performed on Loupe Cell Browser to detect cellular composition and cell type identification using canonical gene expression profiles.

#### Normalization of Samples for Each Patient and Clustering

We processed the unique matrix of gene counts versus cells obtained from normalized aggregate data for each patient using the Seurat (version 2.3.4) R (version 1.1.463) Package (*28*). As a QC step, we first filtered out genes detected in less than three cells and cells that had fewer than 200 genes with read counts greater than zero. Based on the initial data analysis and visual inspection of our dataset, we further filtered cells on percentage of mitochondrial genes and UMI (we chose gene counts over 5,000 and mitochondrial % counts > 0.15) and regressed out the effects of variation in UMI counts and percent mitochondrial counts for those cells remaining. After this step, 24,904 single cells and 18,398 genes in total remained and were included in downstream analyses. Global-scaling library size normalization method was performed in Seurat on the filtered matrix to obtain the normalized count. To focus on more biologically meaningful variation, we used a subset of highly variable genes to perform unsupervised clustering. To identify highly variable genes, the ‘*FindVariableGenes’* function in the Seurat package was used and genes whose log-mean was between 0.0125 and 3.0 and whose dispersion was above 0.5 were identified, resulting in 1,587 highly variable genes. We performed linear dimensionality reduction on the scaled data on these highly variable genes wherein both cells and genes are ordered according to the PCA scores. Statistically significant first 20 principal components were used in downstream clustering analysis determined by implementing a resampling test inspired by the jackstraw procedure (*59*). To partition the cellular distance matrix into clusters, the graph based *FindClusters* function was implemented setting the ‘*dim.use’* parameter equal to 20 and resolution to 1.2, leaving the other parameters as default. To visualize the clusters, cells within the graph-based clusters were localized on the t-SNE plot using *RunTSNE* function using the same principal components as above. Positive and negative differentially expressed genes for single cluster, compared to all other cells was determined using *FindAllMarkers* function on Seurat.

#### Batch-effect Removal Across Individual Patients

Common sources of variation across all the three patients were identified and corrected using canonical correlation analysis (CCA), a method for combining multiple datasets in the Seurat R package (*60*). The union of the top 2,000 genes with the highest dispersion (var/mean) from all three patients was identified to implement *RunMultiCCA* function setting the *num.ccs* parameter to 30 which combined the three datasets as vectors projecting each patient’s dataset into the maximally correlated subspaces. The three datasets are aligned using ‘warping’ algorithms which normalize the low-dimensional representations of each dataset. The CCA subspaces were aligned using *AlignSubspace* function setting the *dim.use* parameter to 1:20, leaving the other parameters as default. Single integrated analysis was performed on all cells to visualize the overlapping clusters.

#### Cluster Cell Type Annotation and Gene Signatures

Two approaches were combined to infer cell types for each cluster. First, we directly examined the expression levels of a set of canonical markers for the target cell types. Enrichment of these markers in certain clusters was considered a strong indication of the clusters representing the corresponding cell types. In the second approach, we assigned cell type to each cluster by correlating cluster mean expression to immune cell expression signatures extracted from the human primary cell atlas (HPCA) dataset using SingleR R package (v0.2.0) (*61*). HPCA, a collection of Gene Expression Omnibus (GEO) datasets contained 713 microarray samples classified to 37 main cell types and further annotated to 157 sub-types (*62*). We annotated a total of 10 T cell clusters, 6 myeloid clusters, 3 NK cell and B cell clusters into different subtypes on co-relation with HPCA dataset.

#### Pseudo-time Trajectory Analysis

To construct cell trajectory manifolds and pseudo-time estimates cells were re-processed using the Monocle R package (v2.8.0) utilizing the reverse graph embedding machine learning algorithm (*63*). The data was normalized for library size using the *estimateSizeFactors* function and negative binomial over-dispersion was estimated for each gene using the *estimateDispersions* function. Dimension reduction was then performed using the DDRTree method on genes selected to have a mean expression value > 0.1 and variance greater than the empirical dispersion. Cells were then represented onto a pseudo-time trajectory using the *orderCells* function (*41*). To identify genes which changed steadily along the identified trajectory, we performed a likelihood ratio test for a negative binomial model using the *differentialGeneTest* function and identified all genes that were significant at a strict 0.01 p-value cut-off after multiple hypothesis correction.

#### Single-sample Gene Set Enrichment Analysis

We carried out ssGSEA (*64*) on the Hallmark gene set library using the GenePattern module *ssGSEA Projection* (v4) (www.genepattern.org) and *SingleR* R package (v0.2.0) (*61*). We used *pheatmap* R package (v1.0.12) for visualization and making heatmaps.

#### Cell Cycle Regression and Re-clustering

To identify different immune cell subpopulations by cellular identity and differentiation states, we extracted all single-cells classified in our first unsupervised clustering analysis and regressed cell cycle activity using the Seurat R Package, as previously described (*65*). The tSNE method was used following the exact steps as described above, testing possible clustering solutions using approaches similar to those described previously.

#### Preprocessing of TCR Sequencing Data and Initial Analysis

The TCR sequencing data from each patient tumor and peripheral blood was preprocessed separately using Cell Ranger 2.1.1, available from 10X Genomics (www.support.10xgenomics.com/single-cell-vdj/software/pipelines/latest/using/vdj)

#### Downstream Analysis of Paired Single-cell RNA-seq and TCR Sequencing Data

The vloupe output file from Cell Ranger VDJ for each sample was analyzed in the Loupe V(D)J Browser, using the import VDJ function for calculating the frequency of shared VDJ clonotypes and CDR3 amino acid sequence between tumor and peripheral blood sample of each patient across all three patients. Further visualization of TCR sequencing data between clusters of individual patients utilized the R packages: dplyr (v0.8.01), reshape2 (v1.4.3), circlize (v0.4.6), ggplot2 (v0.8.2). Individual cell barcodes were assigned as CD4 T cells or CD8 T cells based on the cluster cell type annotations. The productive TCRs for the A and B chain were isolated, excluding multiple chains per barcode. Individual clonotypes between clusters were calculated on per patient, the mean of the shared clonotypes for each cluster was then calculated and displayed in bar graphs. Chord diagram visualizations were modified based on the presumed direction of contribution, i.e., naïve/peripheral blood clusters contribute to tumor-infiltrating immune clusters. Self-contribution of each cluster was removed from the chord diagrams to assist with visualization.

#### Statistical Analysis

Statistical Analyses were performed in R (v3.5.1). Two-sample significance testing utilized Welch’s T test, with significance testing for more than three samples utilizing one-way analysis of variance (ANOVA) with Tukey honest significance determination for correcting multiple comparisons. Differential gene expression for single-cell RNA-seq data was performed in the Seurat R package using the Wilcoxon rank sum test with p-values adjusted using the Bonferroni correction method (*59, 60*). Differential gene expression between branches of the cell trajectory manifold was performed in the Monocle R package using likelihood ratio testing (*63*). Visualization of individual differentially expressed genes across clusters utilized Seurat R package for heatmaps and Monocle R package for violin plots.

## Supplementary Materials

Fig. S1. Distinct cellular identity of immune cells in ccRCC for individual patients.

Fig. S2. CD8+ T cell state heterogeneity and associated gene programs.

Fig. S3. CD4+ T cell state heterogeneity and associated gene programs.

Fig. S4. Abundance and Shared TCR Clonotypes.

Fig. S5. TCR Clonotypes in one T cell in a single patient across all individuals.

Fig. S6. Myeloid cell state heterogeneity and associated gene programs

Table S1. Final cell annotation.

Table S2. CD8 cluster markers.

Table S3. CD8 cluster_0 markers.

Table S4. CD8 cluster_3 and 10 markers.

Table S5. CD8 cluster_6 markers.

Table S6. CD4 cluster markers.

Table S7. CD4 tumor cluster markers.

Table S8. CD4 blood cluster markers.

Table S9. CD4 Treg tumor versus blood markers.

Table S10. CD4 Treg cluster markers.

Table S11. Myeloid cluster markers.

Table S12. Monocytes cluster markers.

Table S13. TAM cluster markers.

Table S14. DC tumor versus blood markers.

Table S15. DC cluster markers.

## Acknowledgments

We thank Michael Knudson, Rita Sigmund, Joe Galbraith, Janice Cook-Granroth, Bethany Kilburg and Celeste Charchalac from University of Iowa Carver College of Medicine, Tissue Procurement Core (TPC) and Genito-Urologic Tissue Repository (GUMER) for receiving biological samples and clinical data. We thank Justin Fishbaugh, Heath Vignes and Michael Shey from the University of Iowa Flow Cytometry Facility. We thank Kevin Knudtson, Mary Boes, Garry Hauser and Mari Eyestone from the Iowa Institute of Human Genetics (IIHG) Genomics Division for planning and assisting use of Next Gen Sequencing (NGS) platforms, Diana Kolb from the IIHG Bioinformatics Division and the University of Iowa High Performance Computing (HPC) facility.

## Author contributions

AV, YZ, and WZ conceived the study. Clinical tissue specimens were procured by KN and YZ. AV and WZ developed experimental methods. AV performed the experiments for next gen sequencing data generation and data processing with conceptual input from IIHG. AV, NB, MC and PV analyzed the data with conceptual input from RWJ, AS and the other authors. AV wrote the manuscript under supervision of AS, RWJ, WZ and YZ. All authors have carefully read the manuscript.

## Data and materials availability

Quantified gene expression counts and V(D)J T cell receptor sequences for single-cell RNA sequencing are available at the Gene Expression Omnibus (GEO) at <u>GSE121638</u>.

## Supplementary Figure Legends

**Fig. S1.**
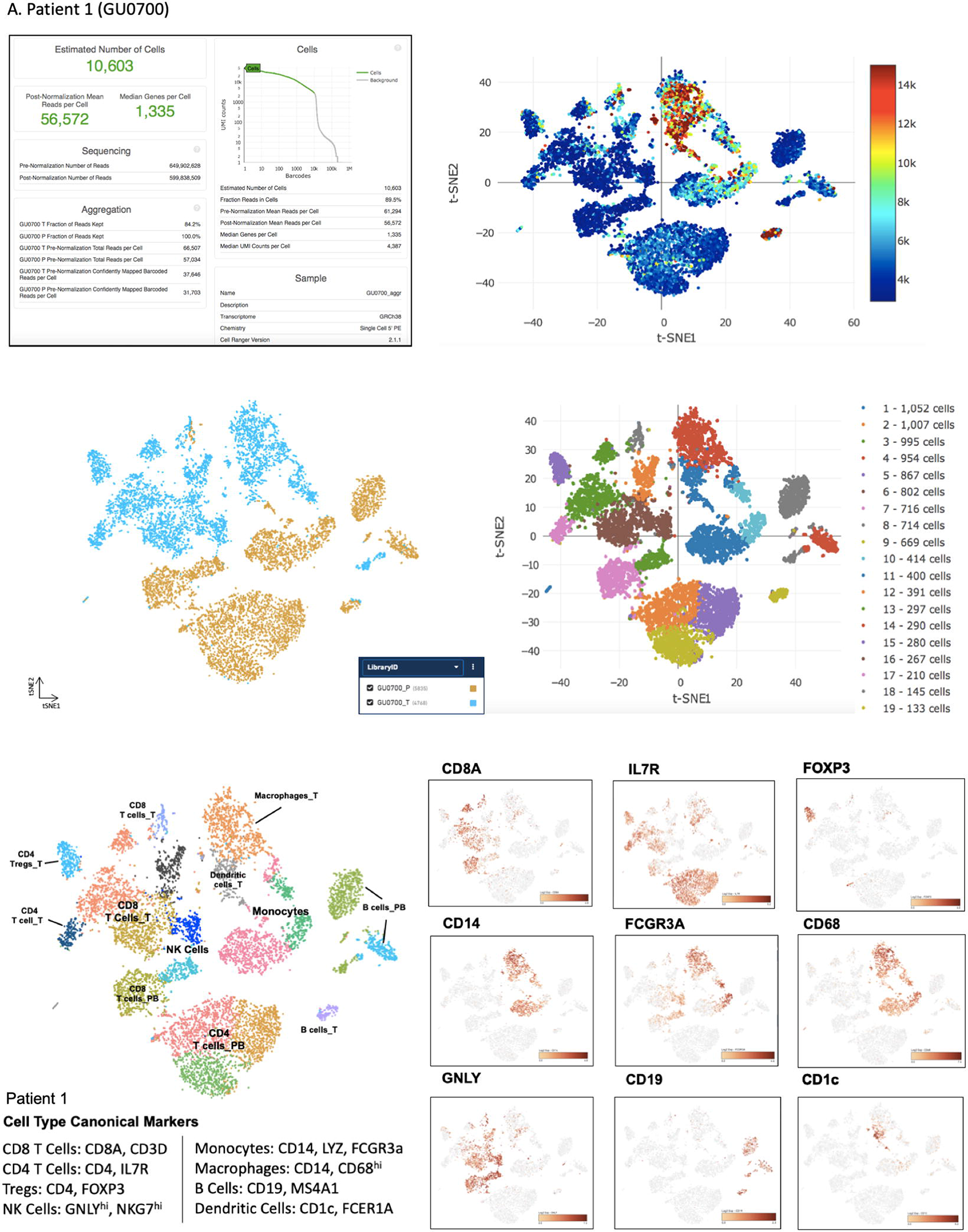
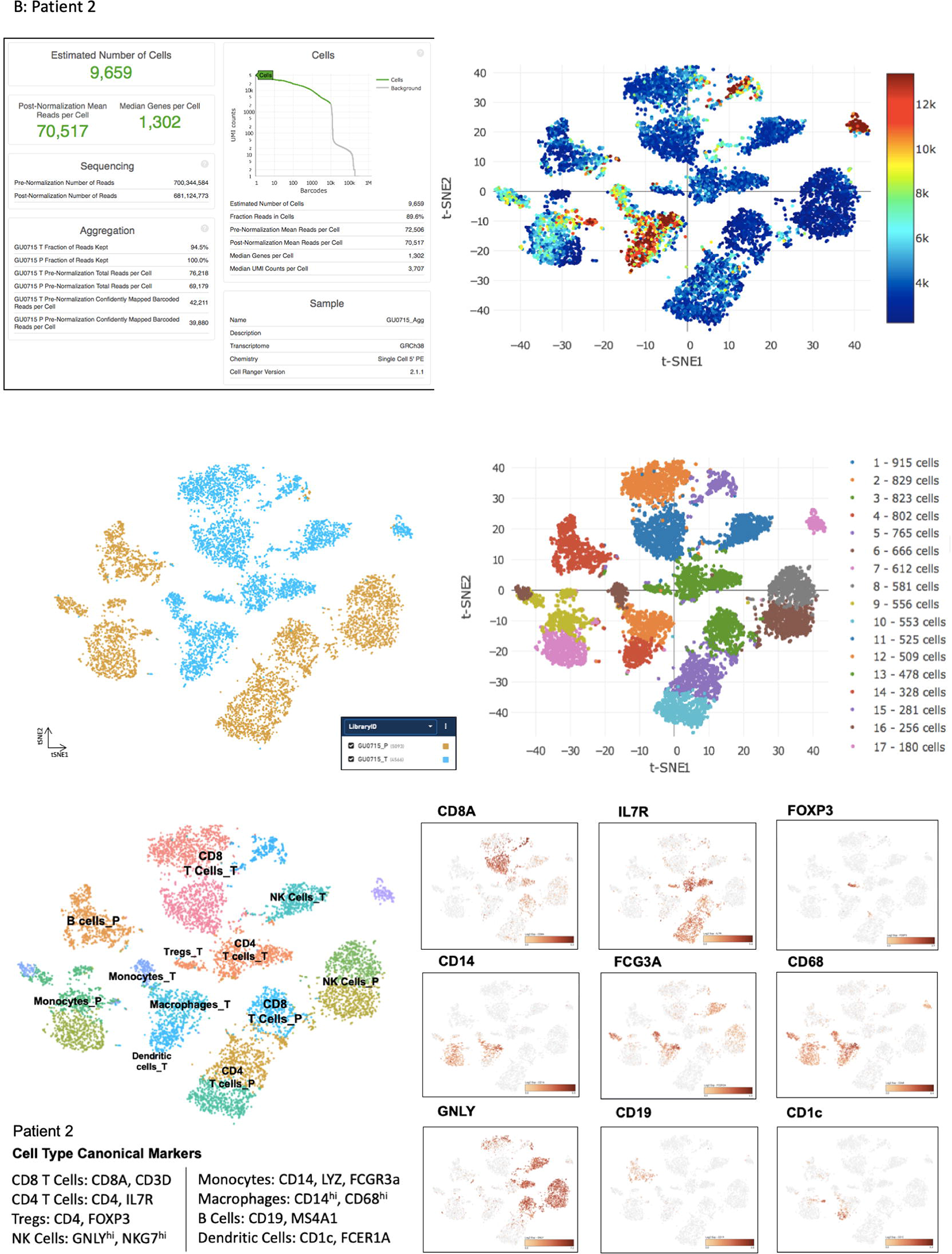
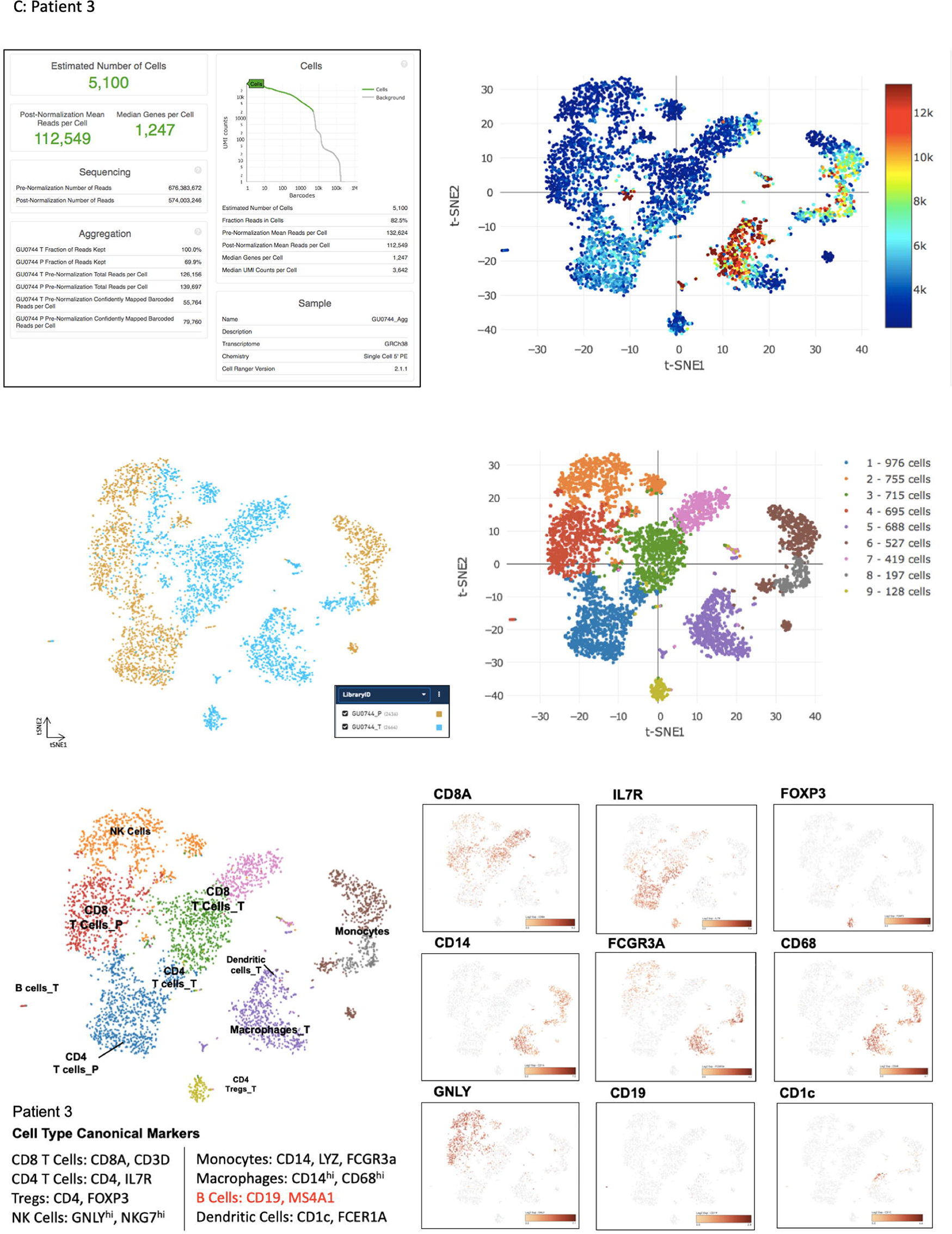
Single-Cell RNA Sequencing Results of All Three ccRCC Patients (S1A, S1B and S1C) from Cell Ranger, 10X software for Initial Data Analysis: The top left panel shows barcode-rank plot. The Y-axis is the number of UMI counts mapped to each barcode and the x-axis is the number of barcodes below that value. A steep drop-off is indicative of good separation between the cell-associated barcodes and the barcodes associated with empty partitions. Top right plot shows tSNE UMI plot. Each point in the map represents an immune cell. Cells with greater UMI likely have higher RNA content than cells with fewer UMI counts. The axes correspond to the 2-dimensional embedding produced by the tSNE algorithm. In this space, pairs of cells that are close to each other have more similar gene expression profiles than cells that are distant from each other. The middle two panels show clustering analysis to illustrate library ID (Tumor versus Peripheral Blood) and basic cell and number of cells in each cluster subsets. The bottom panel shows expression of significant canonical immune genes that are expressed highly within clusters, relative to entire datasets to annotate cell types.

**Fig. S2.**
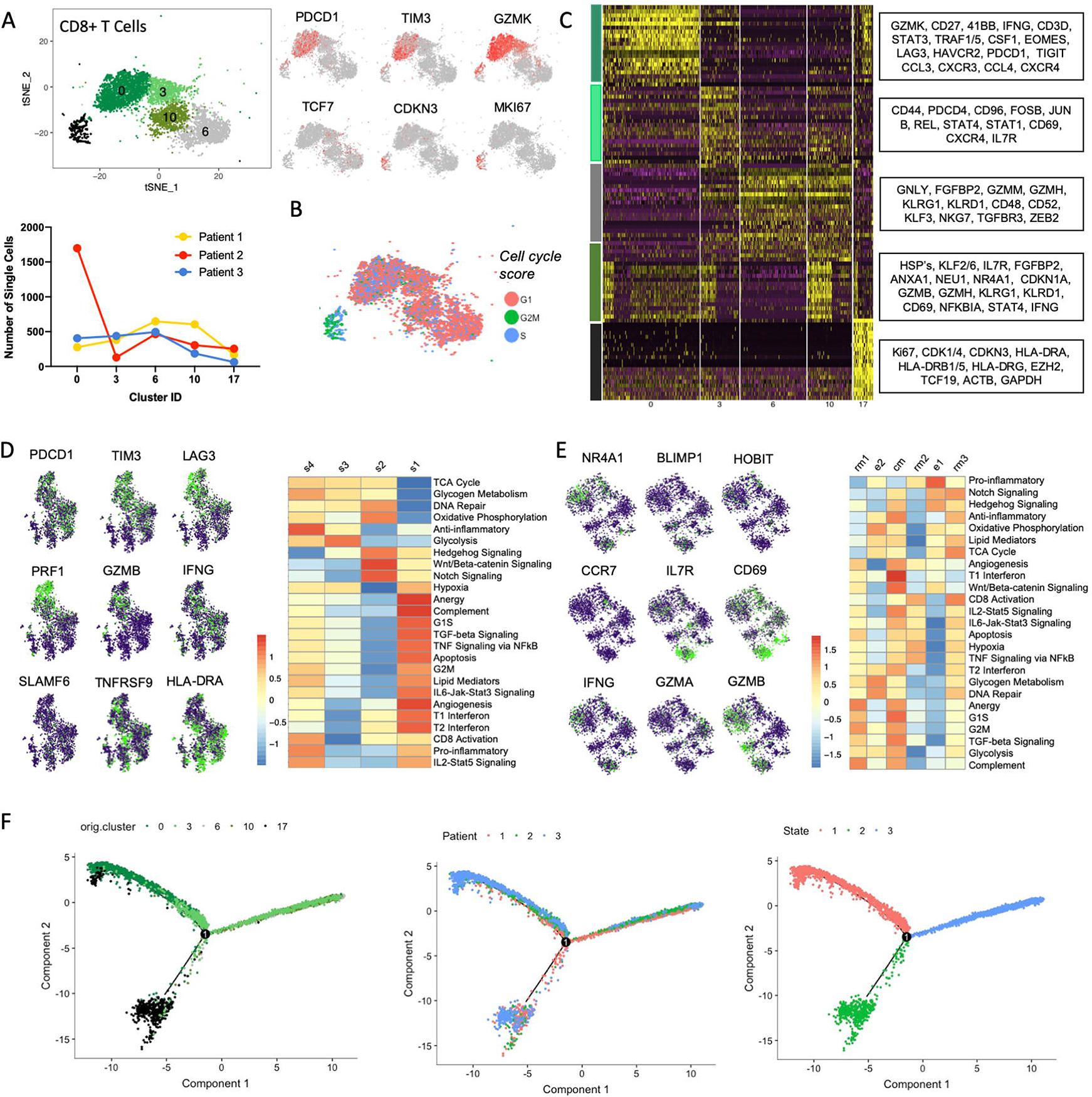
CD8+ T Cell State Heterogeneity and Associated Gene Programs: A tSNE representation of the 6,529 CD8+ T cells and its distribution across all three ccRCC tumor samples along with gene expression values of indicated markers on the tSNE map. (A). A tSNE plot of cell cycle score colored by G1, G2M and S phases (B). The top right panel shows the heatmap expression of the top 20 differentially expressed markers for each CD8+ T cell cluster (see **Table S2**) in (C). Gene expression values of the key indicated markers on the tSNE map of cell cycle regressed CD8_0 cluster and CD8_3 with CD8_10 cluster sub-populations with heatmap representation for ssGSEA enrichment z-scores is shown in (D) and (E) respectively. Pseudotime trajectory plot as in (**Fig 3F**) but this time coloring the cells by original cluster annotation in (A), by patient and by state.

**Fig. S3.**
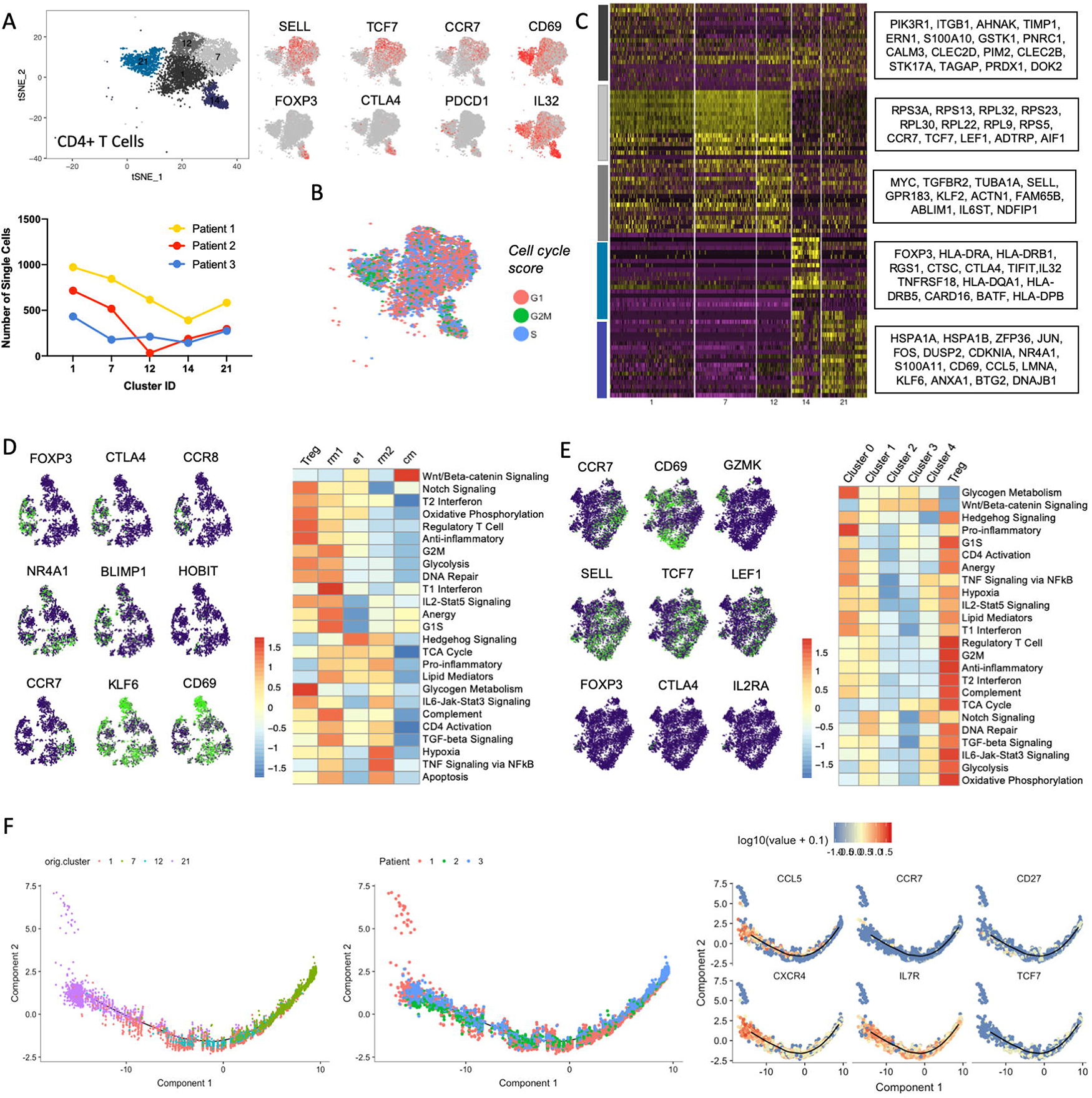
CD4+ T Cell State Heterogeneity and Associated Gene Programs: A tSNE representation of the 6,389 CD4+ T cells and its distribution across all three ccRCC tumor samples along with gene expression values of indicated markers on the tSNE map. (A). A tSNE plot of cell cycle score colored by G1, G2M and S phases (B). The top right panel shows the heatmap expression of the top 20 differentially expressed markers for each CD4+ T cell cluster (see **Table S3**) in (C). Gene expression values of the key indicated markers on the tSNE map of cell cycle regressed tumoral and peripheral cluster sub-populations with heatmap representation for ssGSEA enrichment z-scores is shown in (D) and (E) respectively. Pseudotime trajectory plot of clusters in (A) coloring the cells by original cluster annotation, by patient and key markers.

**Fig. S4.**
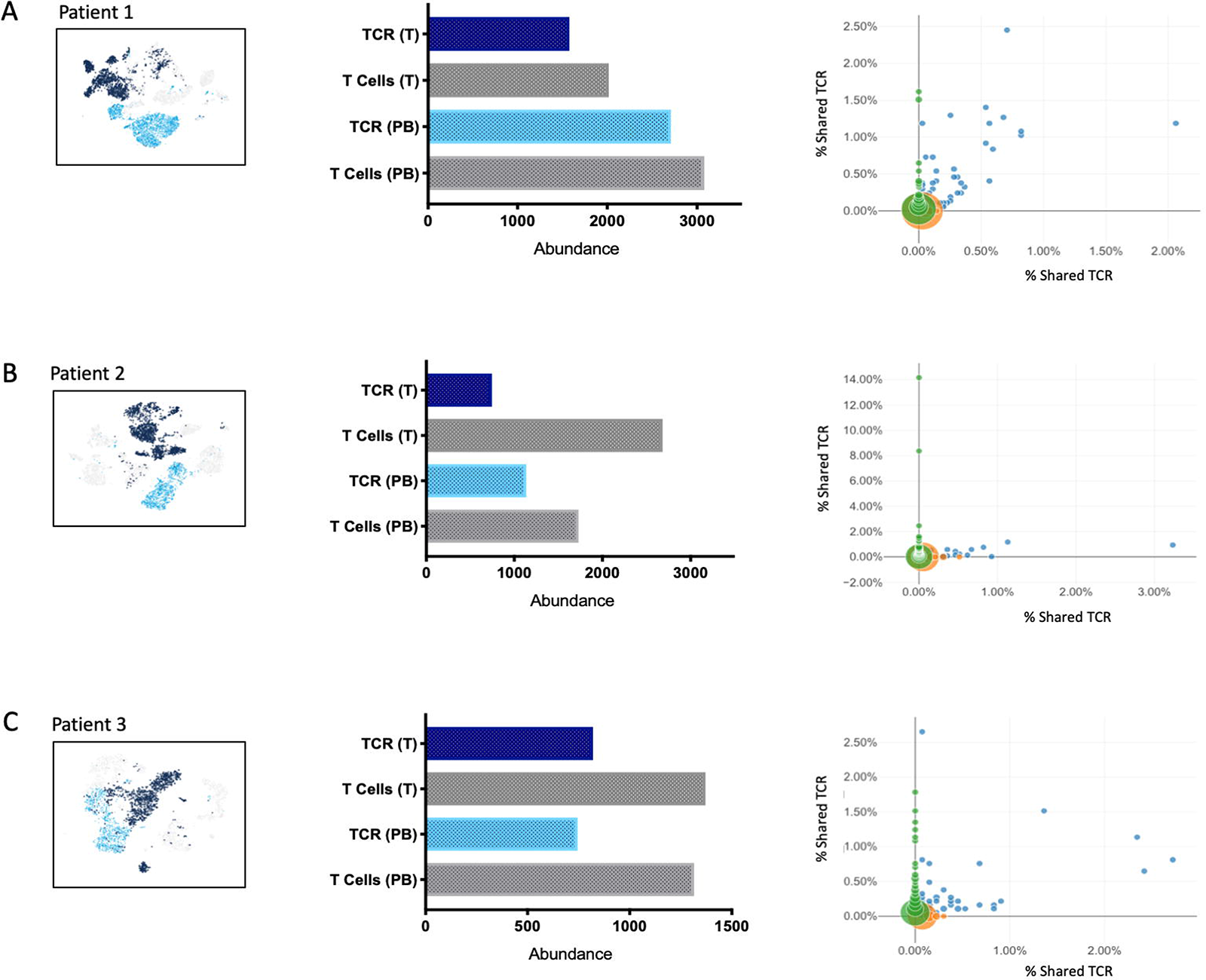
TCR abundance and Shared TCR Clonotypes across Each Patient: A tSNE plot representation of TCR clonotypes in tumor (T) and peripheral blood along with TCR and T cell abundance is shown in left panel and right panel shows shared percentage TCR between the two tissue types in Patient 1 (A), Patient 2 (B) and Patient 3 (C).

**Fig. S5.**
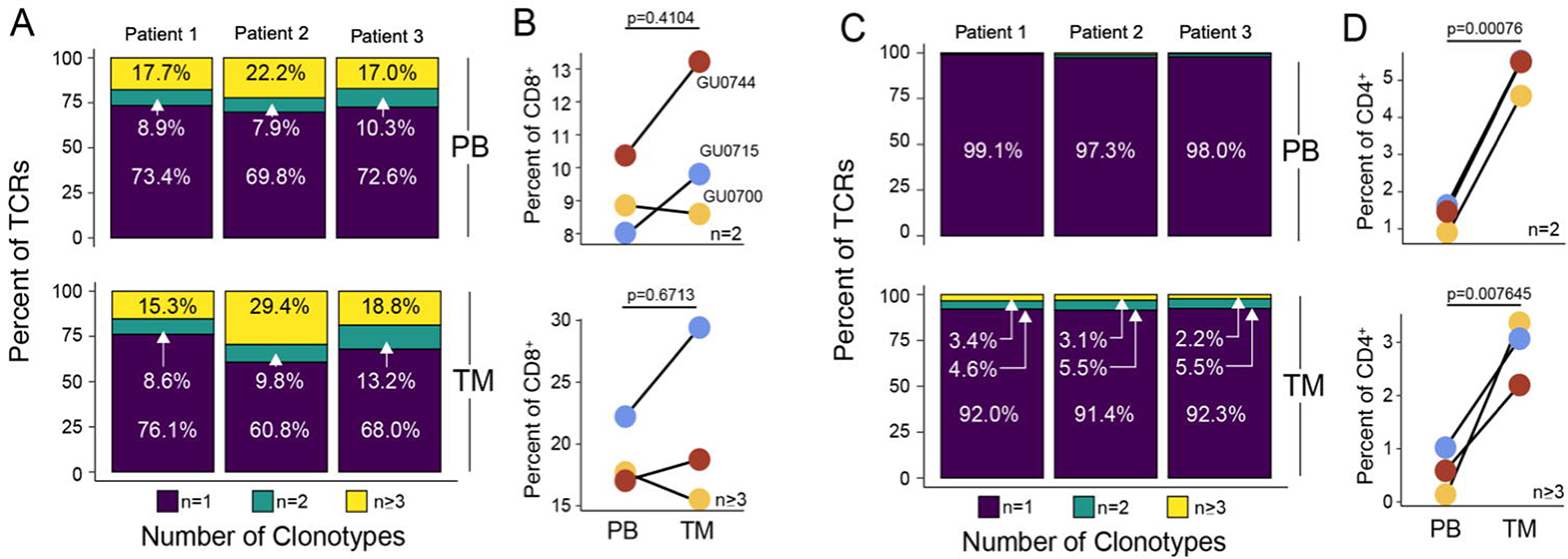
TCR Clonotypes in One T Cell in a Single Patient across All Individuals: Distribution of single, double and three of more T cell clonotypes observed across CD8+ T cell TCRs is shown in (A), (B) and CD4+ T cell TCRs shown in (C) and (D)

**Fig. S6.**
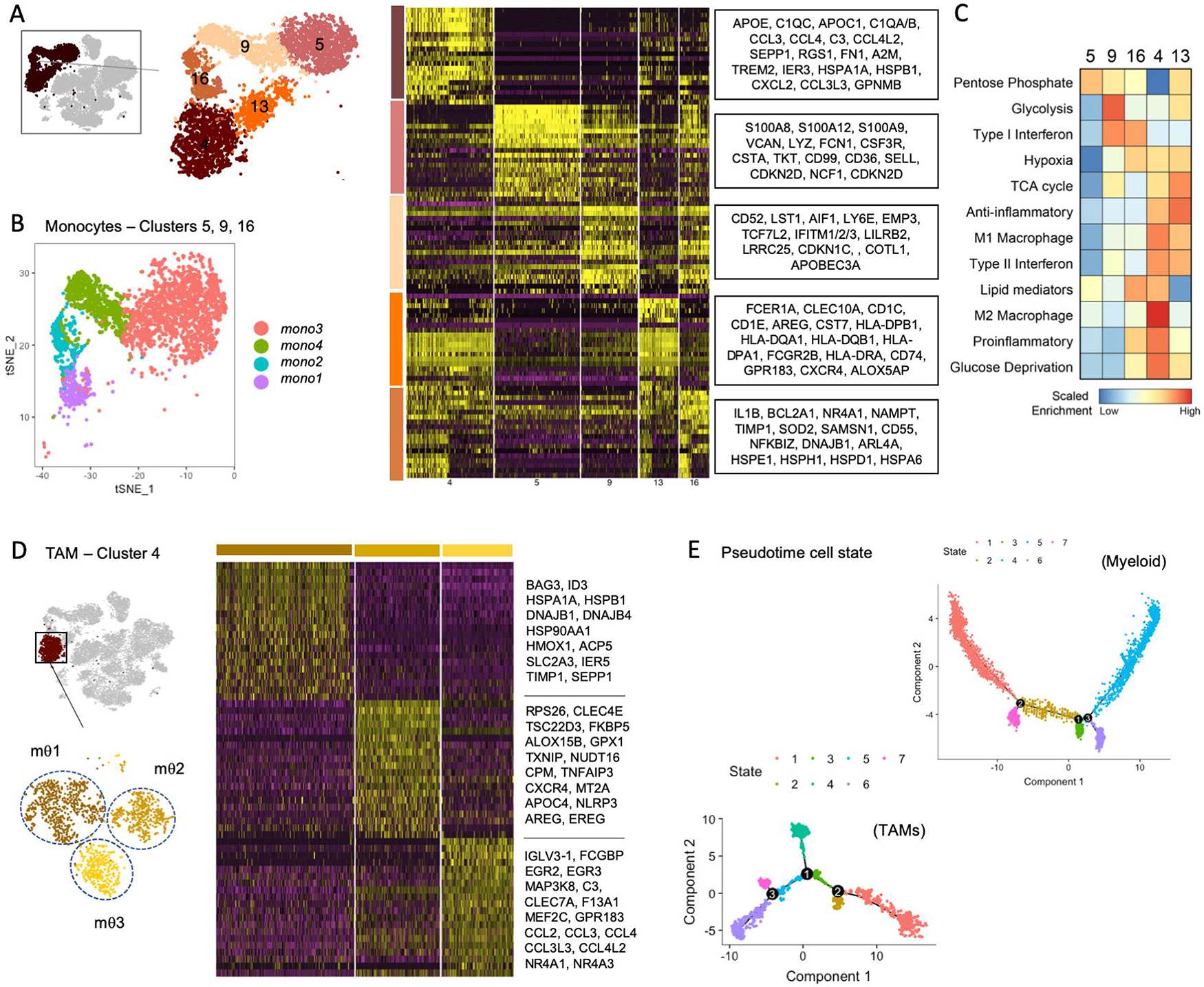
Myeloid Cell State Heterogeneity and Associated Gene Programs: A tSNE representation of 5,940 myeloid cells and heatmap expression of the top 20 differentially expressed markers for each myeloid cell cluster (A). tSNE plot of classical and non-classical monocytes subsets in tumor-infiltrating myeloid cells versus peripheral blood is shown in (B). Heatmap representation for ssGSEA enrichment z-scores is shown in (C) for the myeloid clusters in (A). tSNE plot of tumor-infiltrating macrophage subsets heatmap expression of the top 20 differentially expressed markers for each cell states are shown in (D) along with pseudotime trajectory plot of myeloid clusters (top panel) and TAMs (bottom panel) in (E).

